# PGR5 is required for efficient Q cycle in the cytochrome *b*_6_*f* complex during cyclic electron flow

**DOI:** 10.1101/854489

**Authors:** Felix Buchert, Laura Mosebach, Philipp Gäbelein, Michael Hippler

## Abstract

Proton Gradient Regulation 5 (PGR5) is involved in the control of photosynthetic electron transfer but its mechanistic role is not yet clear. Several models have been proposed to explain phenotypes such as a diminished steady state proton motive force (*pmf*) and increased photodamage of photosystem I (PSI). Playing a regulatory role in cyclic electron flow (CEF) around PSI, PGR5 contributes indirectly to PSI protection by enhancing photosynthetic control, which is a pH-dependent downregulation of electron transfer at the cytochrome *b*_6_*f* complex (*b*_6_*f*). Here, we re-evaluated the role of PGR5 in the green alga *Chlamydomonas reinhardtii* and conclude that *pgr5* possesses a dysfunctional *b*_6_*f*. Our data indicate that the *b*_6_*f* low-potential chain redox activity likely operated in two distinct modes – via the canonical Q cycle during linear electron flow and via an alternative Q cycle during CEF, attributing a ferredoxin-plastoquinone reductase activity to the *b*_6_*f*. The latter mode allowed efficient oxidation of the low-potential chain in the WT *b*_6_*f*. A switch between the two Q cycle modes was dependent of PGR5 and relied on unknown stromal electron carrier(s), which were a general requirement for *b*_6_*f* activity. In CEF-favouring conditions the electron transfer bottleneck in *pgr5* was the *b*_6_*f* and insufficient flexibility in the low-potential chain redox tuning might account for the mutant *pmf* phenotype and the secondary consequences. Models of our findings are discussed.

## Introduction

In linear electron flow (LEF), the two photosystems (PSII and PSI) act in series to ultimately reduce NADP^+^ via the enzyme ferredoxin (Fd)-NADP(H) oxidoreductase (FNR). The cytochrome *b*_*6*_*f* complex (*b*_*6*_*f*) functionally interconnects the two photosystems (reviewed in 1), accepting electrons from plastoquinol (PQH_2_) and donating electrons to plastocyanin (PC). Functional *b*_*6*_*f* occurs as a homodimer, each monomer consisting of four major subunits (cytochrome *b*_*6*_, subunit-IV, cytochrome *f* (cyt.*f*), and the Rieske iron sulphur protein (ISP)), as well as four minor subunits (PetG, L, M and N). In addition, each monomer includes six cofactors: two *b*-hemes (*b*_l_ and *b*_h_), two *c*-hemes (cyt.*f* and *c*_i_), one chlorophyll *a* and one β-carotene. Light indirectly induces *b*_*6*_*f* turnover: Upon oxidation by the primary PSI electron donor P700, PC extracts one electron from cyt.*f*, which is re-reduced by the Rieske ISP. The positively charged Rieske FeS domain moves towards the lumenal Qo-site, where an electron flow bifurcation occurs: PQH_2_ donates one electron to the Rieske ISP (part of the high-potential chain with a midpoint potential E_m_ = 300-350 mV) and a second electron to *b*_l_ (low-potential chain; E_m_ = -130 mV). PQ is re-reduced at the stromal Qi-site via *b*_h_ (E_m_ = -35 mV) and/or *c*_i_ (E_m_ = 100 mV, flexible as described below). Via the canonical Q cycle, the production of one PQH_2_ at Qi requires the oxidation of two PQH_2_ at Qo (supplementary Figure S1). The spatial proximity between *b*_h_ and *c*_i_ suggests electron sharing between the two and the presence of a membrane potential (ΔΨ) promotes the shared electron to rest on *b*_h_^red^/*c*_i_^ox^ (2). Furthermore, presence of a ΔΨ is a general prerequisite for efficient *b*-heme oxidation (3) but it remains enigmatic by which mechanism the *b*_6_*f* senses the ΔΨ. Upon a single Qo-site turnover (Figure S1), the heme-*c*_i_ functions as terminal low-potential chain redox carrier (4). Accordingly, the *b*_*6*_*f* low-potential chain harbours *b*_l_^ox^/*b*_h_^ox^ /*c*_i_^red^ after the first, and *b*_l_^ox^/*b*_h_^ox^ /*c*_i_^red^ after the second Qo-site turnover that is associated with PQH_2_ formation at Qi. The (semi-)PQ in the Qi-site receives the electrons from *c*_i_^red^ and/or *b*_h_^red^ in the presence of ΔΨ, probably in a concerted and closely spaced process. It is of note that heme-*c*_i_ is unique since it lacks an amino acid axial ligand and thus might ligate with the semiquinone analogue NQNO (5, 6) which downshifted the heme-*c*_i_ midpoint potential from 100 mV to approximately -150 mV (4). Furthermore, heme-*c*_i_ was proposed to engage in a Qi-site gating function (7): by ligating tightly in the oxidised state either with the phenyl group of F40 in subunit-IV (closed Qi-site), or, after transient heme-*c*_i_ reduction, with (semi-)PQ. A recent cryo-EM structure of the spinach *b*_6_*f* complex contained the native PQ ligated to heme-*c*_i_ (8). Following ligation-associated midpoint potential downshift of heme-*c*_i_ (4), it is not clarified yet whether heme-*b*_h_ or the quinone is reduced by heme-*c*_i_. Since not more than half the *b*-heme population is reduced per Qo-site turnover in uninhibited complex (9, 10, 11 and references therein), occurrence of *b*_l_^red^/*b*_h_^red^ is unlikely. Part of failing to detect *b*_l_^red^/*b*_h_^red^ might be that, in the presence of *b*_h_^red^ (during a second canonical Q cycle turnover), oxidant-induced reduction of *b*_l_^ox^ provides the strong reducing redox potential in the low-potential chain that is necessary to inject the first electron in the quinone-*c*_i_^ox^ ensemble (7), so that heme-*c*_i_ would force the quasi-concerted PQ reduction (6, 12). The deprotonation of PQH_2_ at Qo and the protonation of PQ at Qi couple electron transfer to proton translocation into the thylakoid lumen. The resulting transmembrane electrochemical proton gradient (*pmf*) fuels ATP synthesis via the chloroplast ATP-synthase. Besides LEF, which produces both NADPH_2_ and ATP, diverse auxiliary electron flow pathways, including cyclic electron flow (CEF) around PSI, contribute to the *pmf* and thereby equilibrate the NADPH_2_ to ATP output ratio of the light reactions (reviewed in 13). In addition, the *pmf* plays an integral photoprotective role, since the chemical component (ΔpH) induces energy-dependent quenching (qE) and modulates the rate-limiting, pH-dependent oxidation of PQH_2_ at the Qo site, which is termed photosynthetic control (14, reviewed in 15). Hence, CEF creates a regulatory feedback loop linking the stromal redox poise to the efficiency of light harvesting and the rate of electron transfer.

PGR5 (proton gradient regulation 5) has been first identified in *Arabidopsis thaliana* as component being involved in the regulation of the *pmf* via CEF (16). The corresponding knockout mutant in *C. reinhardtii* features multi-faceted phenotypes resembling its vascular plant counterpart (17, 18): The algal *pgr5* fails to induce qE-dependent NPQ and is extremely susceptible to PSI photodamage in response to high light (19, 20) as well as fluctuating illumination (21). These defects have been attributed to an impaired acidification of the thylakoid lumen due to compromised Fd-PQ reductase-dependent CEF and a resulting lack of photosynthetic control in response to enhanced stromal redox pressure (19, 22). However, the detailed mechanism of this CEF route is still elusive, as is the molecular role of PGR5. In the past, the association of FNR with the *b*_*6*_*f* (23-25) has been proposed to induce a switch from LEF to CEF: According to this model, FNR would tether reduced Fd in the vicinity of *b*_h_ and *c*_i_, ultimately facilitating PQ reduction via a modified Q cycle that combines lumenal and stromal electrons (26-28). Our previous work showed less stable binding of algal FNR to the thylakoid membrane in the absence of PGR5 (20), suggesting a structural or regulatory contribution of PGR5 to this CEF pathway by influencing the localisation of FNR. By contributing to photosynthetic control and potentially providing the Fd-PQ reductase activity required for CEF, the *b*_*6*_*f* seems to be at the core of the phenotypes the absence of PGR5 produces. Therefore, we spectroscopically reinvestigated the impact of PGR5 on photosynthetic electron transfer in *C. reinhardtii* with a focus on *b*_*6*_*f* functionality by probing the behaviour of the high- and low-potential chain as well as the electrogenic capacity of the photosynthetic machinery. We provide evidence that during CEF a Fd-assisted Q cycle is active which requires PGR5 for sustained *b*_6_*f* function in the light.

## Results

### 1. Alterations in electron transfer in Chlamydomonas *pgr5*

When assessing the function of PGR5 in photosynthetic electron transfer, PSI photodamage in *pgr5* has to be anticipated (19, 20). To re-evaluate the role of PGR5 in electron transfer regulation, we combined several *in vivo* measurement protocols to assess PGR5-dependent electron transfer under low-light autotrophic conditions in *C. reinhardtii*. For this dataset we compared *pgr5* strain with WT, and furthermore investigated a partially PGR5-rescued strain C1 (19). The weak irradiance during growth and measurement provided permissive conditions for the classical *pgr5* phenotype by avoiding photodamage (19, 20). Thus, we obtained a PSI:PSII ratio of 1.13 ± 0.13, 1.18 ± 0.13 and 1.24 ± 0.18 in WT, *pgr5* and complemented C1, respectively (*N* = 3 ± SD). The PSI:PSII ratio was calculated from the electrochromic shift (ECS) amplitude ratio of laser flash-induced charge separations in presence and absence of PSII activity, by using single turnover saturating flashes and adding PSII inhibitors. The actinic light intensity in the measuring cuvette produced an initial electrogenic signal that was characterized by the slope of the ECS intrinsic voltmeter. Similar initial membrane potential (ΔΨ) formation rates were obtained in WT, *pgr5* and C1, respectively, separating 216 ± 23, 193 ± 11 and 185 ± 12 e/s/photosynthetic chain (*N* = 3 ± SD). This actinic background light served to light-adapt cells for at least 30-min in the measuring cuvette. Another parameter that was varied was the stromal redox state which became strongly reduced in anoxic conditions, by depriving oxygen and thus inhibiting mitochondrial respiration (29). Where indicated, oxic control samples were poised with methyl viologen (MV) to abolish PSI acceptor side limitation and inhibit CEF. We functionally analysed both photosystems and the *b*_6_*f* when, except for MV samples, CEF and LEF were operational in light-adapted algae. Before documenting mutant behaviour, we will first describe the WT in more detail in the following three paragraphs.

The experiments were designed so that a steady state, light-adapted system was measured which was briefly perturbed by a saturating light pulse, for multiple turnovers, and then relaxed during several seconds of darkness. We monitored PSI performance in absence of PSII inhibitors first by following the classical Klughammer & Schreiber method (30), in which a saturating ms-pulse is superimposed on actinic background light (Figure 1*A*). A net oxidation of P700 caused negative signals. The redox signals of fully re-reduced P700 after several seconds darkness were set to zero. To achieve full P700 oxidation during the strong pulse, algal PSII was poised with hydroxylamine/DCMU (not shown) which, by serving as reference, allowed to deduce partial oxidation ratios in uninhibited cells (Figure 1*A*). The calculations are shown in Figure 1*B*, and the yield of P700 (YI) was about 0.27 in oxic WT controls. In other words, an additional 27% of light-adapted P700 were photo-oxidisable during the saturating ms-pulse. PSI acceptor side limitation (YNA) in these samples was 0.51 and donor side limitation (YND) was 0.22. Addition of MV abolished YNA and CEF which, despite active PSII, resulted in 57% pre-oxidised P700 in the actinic background light due to YND. Regarding YI, an additional 43% of P700 could get oxidised during the saturating ms-pulse. When oxygen depletion produced strongly reducing conditions, anoxic samples showed an increased YNA in the light and a lower YI fraction of P700. YND was also slightly diminished in anoxic samples.

**Figure 1.**
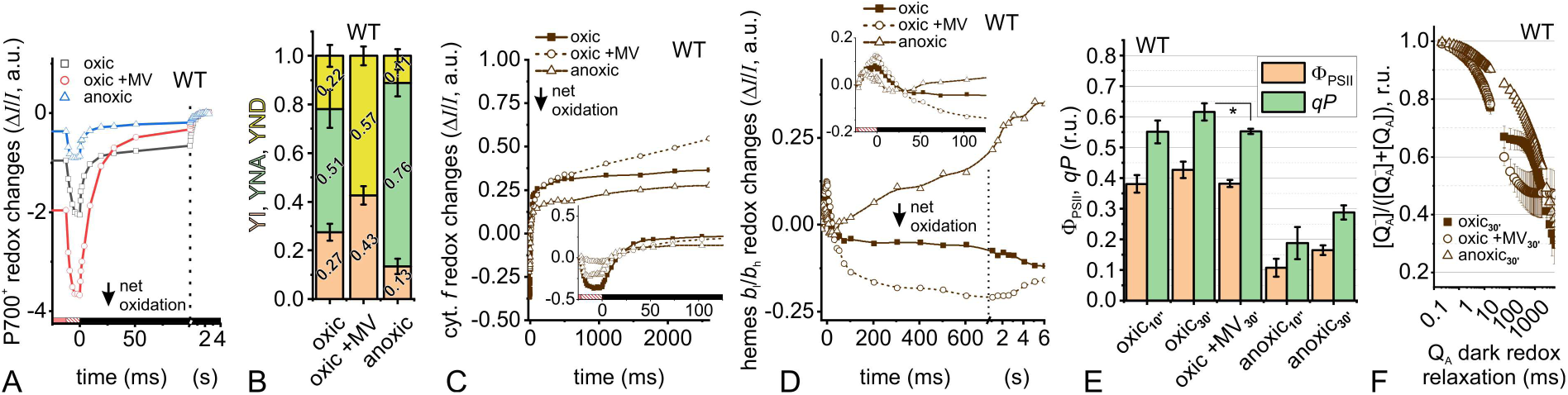
Total electron transfer from PSII to PSI is changed in WT under different cellular redox states. (A) Saturating pulse-induced P700 redox changes in absence and presence of 10 mM methyl viologen (MV) are shown, as well as in anoxic cells. The 12-ms pulse (hatched red box) was applied on light-adapted cells in the steady state (red box), followed by several seconds dark measurements (black box). (B) The different P700 populations were deconvoluted as oxidisable fraction (YI, yield of PSI), non-oxidisable P700 owing to acceptor side limitation (YNA), and pre-oxidised fraction due to donor side limitation (YND). The electron acceptor MV abolished YNA and increased YND despite PSII activity. Anoxic conditions lowered YI and increased YNA. (C) Cytochrome *f* redox changes in a similar light/pulse/dark regime show different pulse-induced oxidation magnitudes (inset) with oxic > anoxic > oxic+MV amplitudes, compared to the steady state reference. MV addition caused no significant cytochrome *f* net oxidation during the pulse. Most of the dark relaxation finished in ∼50-ms, except for slow re-reduction in MV samples. (D) Redox changes of *b*-hemes were comparable during the pulse (inset) and most of the oxidation was finished in ∼50-ms, ∼300-ms, ∼25-ms in oxic, oxic +MV and anoxic samples, respectively. The latter samples showed a slow re-reduction phase with an onset at less than 100-ms dark. (E) Chlorophyll fluorescence-derived quantum yield of PSII (Φ_PSII_) and PSII efficiency factor (*qP*) are shown (*N* = 3 ± SD). The pre-steady state after 10-s illumination and steady state cells after 30-min light adaptation were similar and MV addition significantly lowered *qP* (Welch’s t-test \**P* < 0.05) which, like Φ_PSII_, was also lower in anoxic cells. (F) Following a saturating-pulse in light-adapted cells, redox relaxation of Q_A_^−^ in the dark are shown (*N* = 3 ± SD). The fully reduced Q_A_^−^ pool was re-oxidised in oxic samples by ∼30% in the first ∼50-ms and a following ∼500-ms retardation phase was missing upon MV addition. Q_A_^−^ re-oxidation was slowed down in anoxic cells.

Next, we monitored the *b*_6_*f* redox changes in the same samples. It is of note that the measured signals, especially during the ms-pulse, were sometimes small in amplitude but they were absent in *b*_6_*f*-lacking mutants (supplementary Figure S2). When setting cytochrome *f* (cyt.*f*) redox signals in the background light to zero, the saturating ms-pulse caused a net cyt.*f* oxidation of about -0.4 a.u. in oxic samples (hatched box in insert of Figure 1*C*). In the dark, after the pulse, the fast cyt.*f* net reduction phase had an amplitude of about +0.65 a.u. and finished in ∼50 ms. Both amplitudes of cyt.*f* net oxidation (−0.2 a.u.) and reduction phase (+0.4 a.u.) were smaller in anoxic samples, compared to oxic conditions. The time to reach maximal oxidation during the pulse was shorter in anoxic samples (inset Figure 1*C*), since it was related to the lower YI fraction under these conditions (Figure 1*B*). The cyt.*f* re-reduction kinetics in darkness was faster in these samples as well (inset Figure 1*C*). When MV was present, there was almost no cyt.*f* net oxidation during the pulse. As elaborated further at the end of this section, no distinct fast cyt.*f* reduction phase was observed during 100-ms darkness. When monitoring redox changes of the hemes *b*_l_/*b*_h_ during the saturating ms-pulse in oxic samples (Figure 1*D*), a net reduction of about +0.1 a.u. was observed. Again, redox signals in the background light before the pulse were set to zero. In darkness after the saturating pulse, the hemes *b*_l_/*b*_h_ net oxidation was finished in ∼50-ms in oxic samples and the redox state was slightly more oxidised for several seconds darkness compared to the steady state established in the background light. The hemes *b*_l_/*b*_h_ net reduction amplitude during the ms-pulse was a bit more pronounced in presence of MV. After the pulse, however, there was a significantly larger amplitude of hemes *b*_l_/*b*_h_ net oxidation in the MV samples which transiently reached -0.2 a.u. compared to the pre-equilibrated steady state. The oxidation was finished after ∼300-ms darkness. In anoxic samples during the ms-pulse, hemes *b*_l_/*b*_h_ net reduction amplitude reached a slightly lower plateau earlier. In darkness, hemes *b*_l_/*b*_h_ net oxidation amplitude was small and the phase finished in ∼25-ms. In contrast to oxic samples, a unique hemes *b*_l_/*b*_h_ redox feature was a net reduction phase that started by no later than 100-ms darkness in anoxic WT.

PSII performance was monitored by chlorophyll fluorescence measurements of dark- and light-adapted algae. The maximum quantum yield of PSII, Fv/Fm, was expectedly different in oxic and anoxic samples and was comparable in the strains (supplementary Table S1). When dark-adapted WT was illuminated with the background light for a rather short period of 10-s, the photochemical quantum yield of PSII (Φ_PSII_) as well as the PSII efficiency factor (*qP*) were indifferent from the light-adapted state after 30-min in both oxic and anoxic conditions (Figure 1*E*). In agreement with a previous report (31), *qP* was slightly lower in presence of MV. On the level of Q_A_^−^ re-oxidation after a saturating pulse (32), a multiphasic kinetics was observed in oxic algae during the 6-s of darkness shown in Figure 1*F*. The kinetics of Q_A_^−^ re-oxidation reflect the PQ pool redox state that is in equilibrium with Q_A_ in the dark (32). Starting from an almost fully reduced Q_A_^−^ pool immediately after the pulse (33), the pool became quickly re-oxidized by a fraction of ∼0.3 in the first ∼50-ms of darkness. From there to ∼500-ms darkness, further Q_A_^−^ re-oxidation was absent in oxic samples and a second slow re-oxidation phase followed thereafter. It is of note that the retardation phase between 50-ms and 500-ms darkness, representing downstream electron transfer via the *b*_6_*f* (32), was not established in dark-adapted algae after a short 10-s illumination (supplementary Figure S3). The light adaptation-specific dark kinetics (during progressing *pmf* consumption) link the redox equilibration via the *b*_6_*f* to a reservoir of electrons accumulating downstream of PSI. In MV-treated samples, Q_A_^−^ re-oxidation was identical in the first ∼20-ms darkness compared to oxic controls (Figure 1*F*). The above-mentioned Q_A_^−^ re-oxidation slow down between 50-ms and 500-ms is lacking in MV-treated samples, suggesting a somewhat faster Q_A_^−^ re-oxidation in presence of MV (31) since no electrons could accumulate downstream of PSI. In strongly reduced anoxic cells, Q_A_^−^ re-oxidation was slower during the first ∼500-ms darkness (Figure 1*F*).

When the above experiments were performed in the *pgr5* mutant, clear differences were observed in the electron transfer chain compared to WT: in oxic *pgr5*, a careful look at the kinetics during the ms-pulse showed a faster establishment of P700^+^ plateau (hatched box in Figure 2*A*), which will be explored in more detail at the end of this section. The fractural P700 redox analysis in anoxic conditions revealed that YNA was strongly diminished in *pgr5* and YND appeared to be increased to 0.33 (Figure 2*B*). Figure 2*B* also shows that oxic samples were like WT and MV addition produced slightly less YND in *pgr5*. The net cyt.*f* oxidation amplitude during the pulse was smaller in oxic *pgr5* (−0.2 a.u., Figure 2*C*). In this sample, the reduction phase amplitude was slightly larger (+0.8 a.u.) but the decay kinetics resembled WT, which will also be shown further below. Whereas MV samples were indistinguishable from WT, cyt.*f* redox changes in anoxic *pgr5* differed in several aspects. The net oxidation amplitude was almost non-existent in anoxic cells whereas the reduction amplitude was, unlike the oxic vs. anoxic difference in WT, indistinguishable from oxic *pgr5*. Unlike anoxic WT, fast cyt.*f* reduction kinetics were absent after the pulse and, in fact, slowed down significantly in *pgr5*. The hemes *b*_l_/*b*_h_ redox signals during and after the saturating ms-pulse were like WT with two exceptions in anoxic *pgr5* (Figure 2*D*). The hemes *b*_l_/*b*_h_ oxidation phase after the pulse finished a bit later than in WT and the onset of re-reduction was significantly delayed, which will be quantified further below. Finally, we also observed differences on the level of PSII but this time in the oxic samples. Light-adapted oxic *pgr5* showed higher Φ_PSII_ which, as proposed previously (19), might be the result of higher LEF rates in the mutant (asterisk in Figure 2*E*). In line with this finding, oxic light-adapted *pgr5* samples also showed higher *qP*. When plotting the Q_A_^−^ redox relaxation after full reduction by a saturating pulse (Figure 2*F*), the fast Q_A_^−^ re-oxidation phase up to 20-ms darkness and the slow phase after 500-ms were similar in oxic *pgr5* and WT (cf. Figure 1*F*). However, oxic light-adapted *pgr5* lacked the Q_A_^−^ re-oxidation slow down between 50-ms and 500-ms darkness (Figure 2*F*), which may point to less accumulated electrons downstream of PSI. The latter would be in line with previous studies in Arabidopsis *pgr5* which assigned a disturbed electron flow downstream of PSI, probably on the level of Fd (34, 35). The kinetics in the presence of MV and in anoxic cells were similar, the latter being slightly faster beyond 100-ms darkness in *pgr5*. Establishing the slowdown phase between 50-ms and 500-ms, as a distinguishable feature of the oxic WT and C1 line, required light adaptation for more than 10-s (supplementary Figure S3).

**Figure 2.**
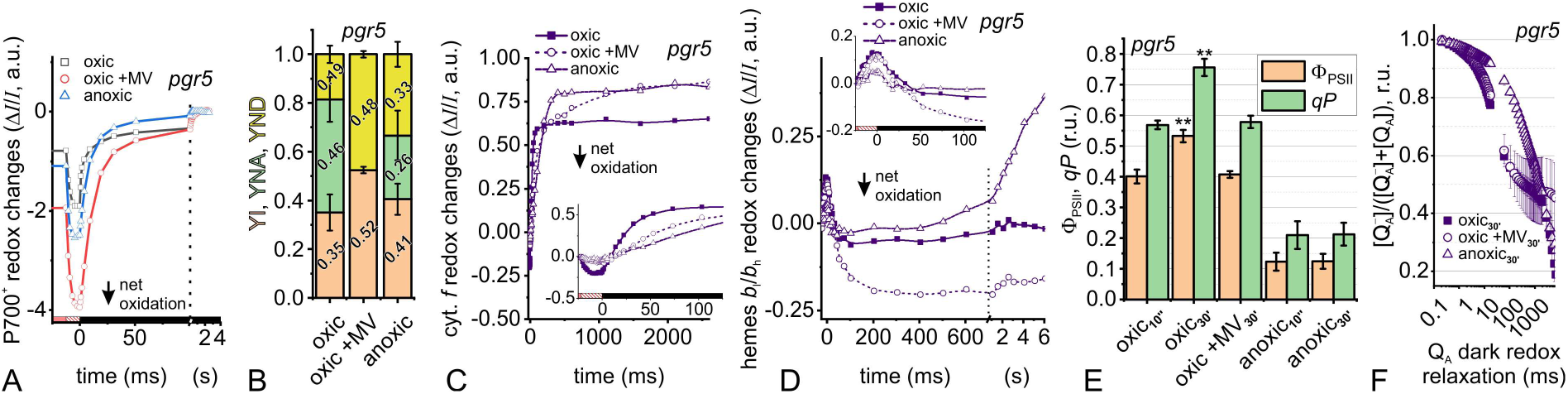
Total electron transfer from PSII to PSI is changed in *pgr5*, most prominently in anoxic conditions. (A) Saturating pulse-induced P700 redox changes in absence and presence of 10 mM methyl viologen (MV) are shown, as well as in anoxic cells. The 12-ms pulse (hatched red box) was applied on light-adapted cells in the steady state (red box), followed by several seconds dark measurements (black box). (B) The different P700 populations were deconvoluted as oxidisable fraction (YI, yield of PSI), non-oxidisable P700 owing to acceptor side limitation (YNA), and pre-oxidised fraction due to donor side limitation (YND). The electron acceptor MV abolished YNA and increased YND despite PSII activity. Unlike WT (cf. Figure 1), anoxic *pgr5* did not produce lower YI and maintained a low YNA fraction. YND was slightly increased (C) Cytochrome *f* redox changes in a similar light/pulse/dark regime show different pulse-induced oxidation magnitudes (inset) which, with oxic > anoxic = oxic+MV amplitudes, compared to the steady state reference. Oxidation amplitudes in oxic controls were smaller than in WT. Mutant kinetics in the presence of MV and in anoxia were similarly slowed down. (D) Redox changes of *b*-hemes were comparable during the pulse (inset) and showed a similar trend as WT with a longer oxidation phase in the presence of MV and slightly shorter phase in anoxia. The latter samples showed a re-reduction phase which started later and was slower than in anoxic WT. (E) Chlorophyll fluorescence-derived quantum yield of PSII (Φ_PSII_) and PSII efficiency factor (*qP*) are shown (*N* = 3 ± SD). The pre-steady state after 10-s illumination and steady state cells after 30-min light adaptation differed in oxic conditions compared to WT (Welch’s t-test \*\**P* < 0.005, cf. Figure 1*E*). This effect was inhibited by MV which yielded as low Φ_PSII_ and *qP* pre-steady state samples without MV. Both parameters were also lower in anoxic cells. (F) Following a saturating-pulse in light-adapted cells, redox relaxation of Q_A_^−^ in the dark are shown (*N* = 3 ± SD). The fully reduced Q_A_^−^ pool was re-oxidised in oxic samples by ∼30% in the first ∼50-ms and, unlike as oxic WT, the ∼500-ms retardation phase was missing in *pgr5* even without MV addition. Q_A_^−^ re-oxidation was slowed down in anoxic cells.

As mentioned above, the rates of P700 oxidation have been determined as well as dark relaxation kinetics on the levels of P700, cyt.*f* and hemes *b*_l_/*b*_h_ (Figure 3). The PGR5-complemented C1 has been analysed (supplementary Figure S4) and is also included. Single-exponential functions were used to determine *k*_*P*-ox_, the P700 oxidation rates during the pulse (Figure 3*A*). When following a previous assumption (19) by ascribing the origin of higher Φ_PSII_ in oxic *pgr5* (Figure 2*E*) to higher LEF rates, the faster P700 oxidation rate during the ms-pulse (Figure 3*A*) would support this hypothesis of faster electron flow via *pgr5* PSI. In the light of undelayed Q_A_^−^ re-oxidation as proxy for missing electron accumulation downstream of PSI in oxic *pgr5* (Figure 2*F*), the faster *k*_*P*-ox_ may be tied to unregulated electron flow between PSI and its sinks (34, 35) which could contribute to diminished photosynthetic control after the saturating pulse. The C1 line was indistinguishable from WT. Since the light intensity during the pulse was saturating, a faster oxidation rate in all anoxic samples could result from a combination of events in these conditions (Figure 3*A*). Higher *k*_*P*-ox_ might reflect a larger PSI antenna size in anoxic cells because of state transitions (36). Since PSI acceptor side limitation was enhanced in the reducing anoxic WT and C1 line (Figure 1*B*, Figure S4*B*), a faster apparent P700 oxidation during the multiple turnover pulse could result from a smaller pool of photoactive PSI. The latter would stem from inefficient charge stabilisation at the PSI acceptor side and the hampered relaxation to the oxidized P700 – the multiple turnover process during the saturating pulse to empty the immediate donor pool of reduced PC and cyt.*f* (30). As elaborated above (Figure 1*B*, Figure 2*B*, Figure S4*B*), PSI donor side limitation was enhanced in the presence of MV which suggests facilitated oxidation of primary and secondary electron donors of P700. It could explain why WT P700 was slightly faster oxidized (Figure 3*A*). However, slightly faster P700 oxidation was not observed in MV-containing *pgr5* and C1.

**Figure 3.**
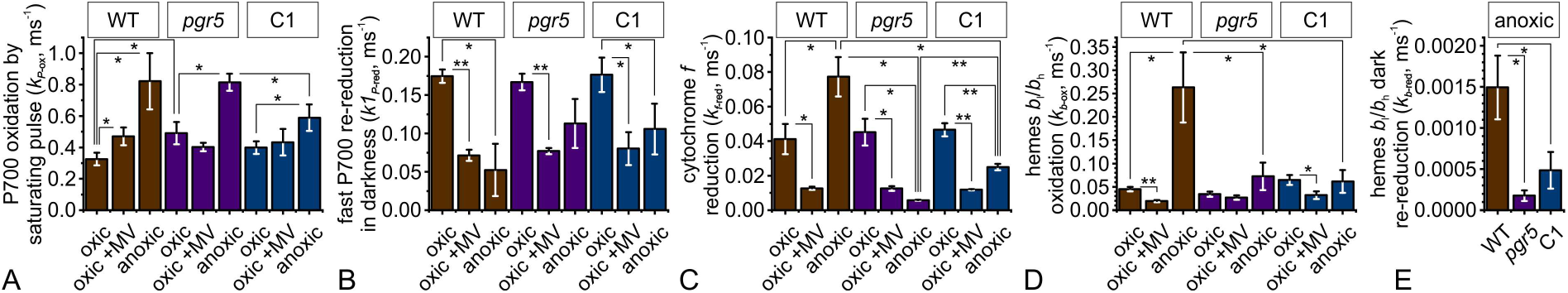
Depending on the conditions and strains, different rates were observed for the redox reactions in P700 and *b*_6_*f* during the saturating pulse and in the dark. Data from Figure 1, Figure 2, and Figure S4 were used for calculations (*N* = 3 ± SD; Welch’s t-test \**P* < 0.05 and \*\**P* < 0.005). As mentioned in the methods, exponential decay functions were used for fitting the data. (A) The pulse-induced P700 oxidation rate, *k*_*P*-ox_, was faster in oxic *pgr5*. Only MV samples in the WT showed faster *k*_*P*-ox_. Anoxic cells also produced faster *k*_*P*-ox_ than oxic controls. (B) The faster of the two P700 re-reduction phases after the pulse yielded *k1*_*P*-red_. This rate was slowed down in MV samples. The anoxic WT and C1 also showed slower *k1*_*P*-red_. (C) Following the pulse, fitting the cytochrome *f* reduction phase yielded *k*_*f*-red_ which was slowed down by MV. Anoxic WT showed faster *k*_*f*-red_, whereas it was slower to a different degree in anoxic *pgr5* and C1. (D) Following the pulse, *b*-heme oxidation rates were calculated as *k*_*b*-ox_ which was slowed down by MV in WT and C1. In anoxic WT *k*_*b*-ox_ was faster compared to the oxic control and the other anoxic strains. (E) The slow *b*-heme re-reduction rate *k*_*b*-red_ in anoxia was fastest in WT.

P700 reduction in the dark occurred in multicomponent decay kinetics (37). During the first few ms after the pulse and especially in reducing conditions, the probability of charge recombination events between P700^+^ and its primary and secondary reduced acceptors increases (38). We therefore disregarded the initial 4 ms of darkness and calculated the first (fast) component, *k1*_*P*-red_, of a 2.5-s two-exponential decay as an estimate for the rate of electron flow that was occurring in the light (Figure 3*B*). We observed in all anoxic cells that there was a slowdown of *k1*_*P*-red_ after the pulse in darkness, to a significant extent in WT and C1. One possibility to explain the P700 reduction slowdown would be enhanced photosynthetic control in these samples. The large build-up of a ΔpH, reported in anoxic algae (39), would slowdown PQH_2_ oxidation at the *b*_6_*f* Qo-site and therefore reduction of PC. All strains showed a smaller *k1*_*P*-red_ in presence of MV (asterisk in Figure 3*B*), which could result from a strongly pre-oxidized pool of primary and secondary donors. Photosynthetic control would slowdown the total electron transfer rate and the PQ pool would become more reduced due to PSII activity. However, a putatively enhanced photosynthetic control contradicts the faster Q_A_^−^ re-oxidation (Figure 1*F*, Figure 2*F*, Figure S4*F*), which is in equilibrium with the PQ pool in the dark (32) and should be slowed down if the latter was in fact more reduced. Later, we will revisit the smaller *k1*_*P*-red_ in presence of MV in the light of *b*_6_*f* performance.

When calculating the reduction rate of cyt.*f* from exponential decay of the kinetics after the pulse (*k*_*f*-red_, Figure 3*C*), oxic samples were identical and significantly slowed down in the presence of MV. Using a multiple turnover protocol, it is important to stress that *k*_*f*-red_ in Figure 3*C* also strongly depended on the oxidation level of the PC pool at the end of the pulse which would have influenced the redox equilibration time between PC and cyt.*f*. Similarly, the PQ pool redox state was also relevant for the calculated rates. When assuming a lake model for light harvesting (40), the PQ pool was, in fact, more reduced in the steady state in anoxic *pgr5* (supplementary Table S1). The PC pool was expected to be oxidized the presence of MV and to answer the question whether reduction of cyt.*f* in Figure 3*C* is truly slowed down, we performed single turnover experiments as well in Results section 3. In addition to the possible impact of the PC pools redox states in Figure 3*C*, there is a *b*_6_*f* intrinsic process, explored in detail further below, which regulates the electrons entering the high-potential chain via a mechanism that depends on electrons exiting the low-potential chain (reviewed in 41).

The latter process was measured from exponential decay of the kinetics after the pulse (*k*_*b*-ox_, Figure 3*D*). MV addition significantly lowered *k*_*b*-ox_ in WT and C1. Oxic *pgr5* controls produced relatively low rates and the inhibitory effect of MV was statistically not significant. Considering the relatively large oxidation amplitudes after the pulse in the MV samples (Figure 1*D*, Figure 2*D*, Figure S4*D*) in combination with lower *k*_*b*-ox_ (Figure 3*D*), it is likely that electrons accumulated in the low-potential chain during illumination which were slowly transferred to PQ at the Qi-site in the dark. According to the above-mentioned coupling concept, the lack of electrons exiting the low-potential chain in presence of MV could result in low *k*_*f*-red_ shown in Figure 3*C*. The anoxic WT produced higher *k*_*b*-ox_ (Figure 3*D*) which, again, would enhance the efficient release of “trapped” high-potential chain electrons (Figure 3*C*). On the contrary, the *pgr5* mutant and the partially complemented C1 strain did not produce higher *k*_*b*-ox_ (Figure 3*D*), thus failing to efficiently reduce cyt.*f* (Figure 3*C*). The C1 line was less severely affected than *pgr5*. We also calculated the slow dark-reduction rates of hemes *b*_l_/*b*_h_ in anoxia from single-exponential fits (*k*_*b*-red_, Figure 3*E*) and the WT, which also showed an earlier onset of this phase, was significantly faster than C1 and *pgr5*. The re-reduction phase in anoxic cells most likely represents re-reduction of heme-*b*_h_, which is accomplished in a few seconds in the dark (42, 43). The origin of the slowdown is unknown, but it could be that delayed hemes *b*_l_/*b*_h_ oxidation in *pgr5*/C1 lowered apparent re-reduction rates.

### 2. Electrogenic capacity of the photosynthetic chain in Chlamydomonas *pgr5*

The data in Figure 1 and Figure 2 indicate that there were alterations in the mutant electron transfer chain. In a similar fashion to the P700 measurements where a superimposed saturating light pulse empties the immediate donor pool of PSI during several turnovers (30), we analysed the charge separation capacity of the photosynthetic apparatus by recording the electrochromic shift (ECS) which is shown in Figure 4 (see supplementary Figure S5 for C1). The ECS, which serves as intrinsic voltmeter (reviewed in 44), was recorded in the background light, during and after the saturating pulse. In the steady state, before the short pulse, the slope of the signal was zero. The additional membrane potential (ΔΨ) that was built up at the very onset of the 22-ms pulse depended on the immediate availability of electrons in the photosynthetic chain and the photochemical yield of both photosystems, which we determined above. The rate of the initial ΔΨ generation, *k*_ini_, was derived from the positive slope during the first 2-ms of the pulse, which is shown as green symbols in WT (inset of Figure 4*A*) and *pgr5* (inset of Figure 4*B*). Oxic samples produced ∼4 additional, stable charge separations during the pulse, also in the presence of MV. Anoxic cells generated less additional ΔΨ during the pulse, and WT was slightly more competent than *pgr5*. During several seconds of darkness when ATP synthase was active, the ΔΨ collapsed to ∼-4 units in oxic and anoxic samples, and to ∼-8 units in MV samples. It is of note that we did not check for the total *pmf* partitioning between ΔpH and ΔΨ. Instead, Figure 4 probed only the ΔΨ which, if it was the dominating *pmf* component, would produce large amplitudes when driving ATP synthesis during 2.5-s darkness. It remains to be tested whether this effect was contributing to the large ECS amplitudes after the pulse in the MV samples. At the end of the pulse a new steady state (zero slope) was established in oxic and anoxic WT only. Upon exhaustion of the immediate donor pool, the ΔΨ generation efficiency at the end of the pulse was derived from a Dark Interval Relaxation Kinetics-based protocol (45). To obtain the progressed ΔΨ production rate at the end of the pulse, *k*_end_, the apparent light-dependent ΔΨ formation (slope of orange symbols in Figure 4*A* and Figure 4*B*) had to be corrected for the ATP synthase-dependent ΔΨ consumption (negative slope in the dark shown by yellow symbols in Figure 4*A* and Figure 4*B*), assuming that the latter was identical just before onset of darkness.

**Figure 4.**
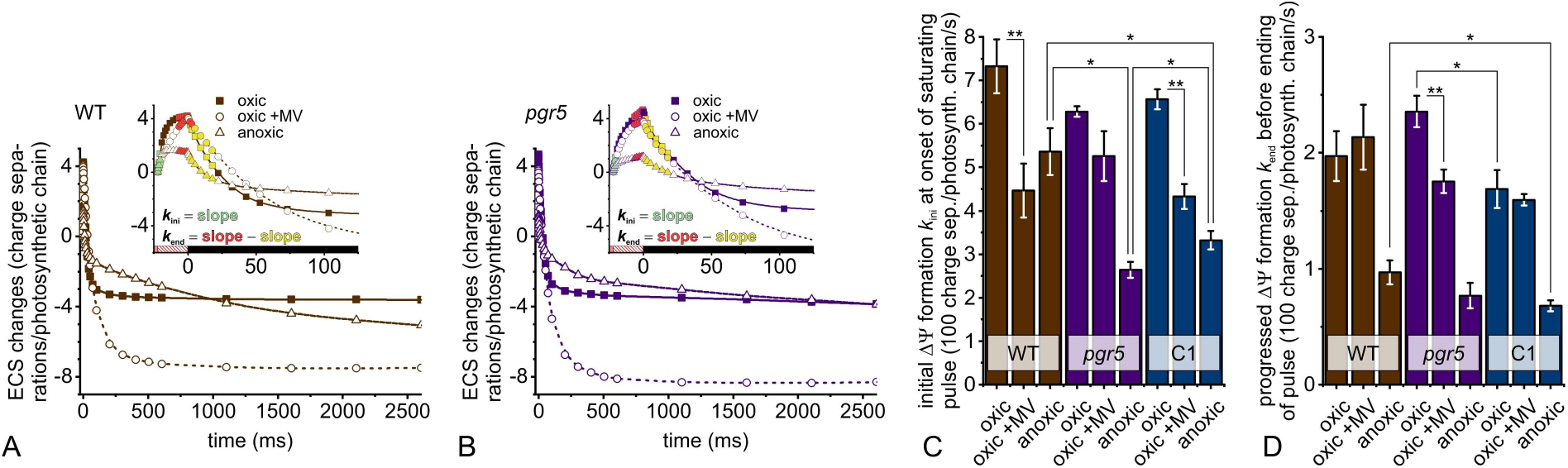
The electrogenic capacity of the photosynthetic electron transfer chain in *pgr5* is compromised under anoxic conditions. The ability to generate and dissipate an electric field is shown by measuring the electrochromic shift, ECS, and signals of samples from Figure 1, Figure 2, and Figure S4 were recorded in steady state light, during a saturating pulse and in darkness. The ECS kinetics for the oxic WT (A) and *pgr5* (B) indicate that the 22-ms pulse led to equilibration of a new membrane potential ΔΨ (for ECS kinetics of C1 refer to supplementary Figure S5). The efficiency to generate a higher ΔΨ level at the onset of the pulse is shown by initial rates in green (inset), yielding *k*_ini_ from the linear slope during the first 2-ms of the pulse. The ΔΨ generation capacity at the end of the pulse (*k*_end_) was corrected with the ΔΨ-consuming activity of the ATP synthase. (C) The *k*_ini_ was similar in oxic strains and slower upon MV addition. A significant MV effect was not observed in *pgr5* which had the lowest *k*_ini_ among the controls. In anoxic cells, *k*_ini_ was highest in WT, followed by C1 and *pgr5*. (D) At the end of the pulse, *k*_end_ was lower in the strains due to exhaustion of electron carriers. There was a difference between oxic *pgr5* and C1. Anoxic *k*_end_ were the same in the strains. (*N* = 3 ± SD; Welch’s t-test \**P* < 0.05 and \*\**P* < 0.005)

The *k*_ini_ values are shown in Figure 4*C* and indicate that in presence of MV, with exception of *pgr5*, the pulse-induced ΔΨ formation was less efficient compared to controls. Nonetheless, the MV effect existed in *pgr5* as well which showed a relatively low *k*_ini_ in oxic controls already. Considering the electrogenic competence associated with *qP*, Φ_PSII_ and YI in the presence of MV (Figure 1, Figure 2 and Figure S4), the observed slowdown of *k*_ini_ (Figure 4*C*) was likely resulting from the poorly functioning *b*_6_*f* (Figure 3*C*-*D*). As will be discussed later, such a situation would imply lower *b*_6_*f*-borne electrogenicity and less immediately oxidisable PC. The enhanced *b*_6_*f* low-potential chain performance in anoxic WT (Figure 3*C*-*E*; in the presence of lower *qP*, Φ_PSII_ and YI) could explain why *k*_ini_ was least diminished among the anoxic strains in Figure 4*C*. Note that, compared to oxic controls, YI was unaffected in anoxic *pgr5*, whereas Φ_PSII_ and *qP* were like in WT. Still, *k*_ini_ and the *b*_6_*f* rates were lowest in anoxic *pgr5*. As expected upon exhaustion of the immediate donor pool, *k*_end_ was diminished compared to *k*_ini_ (Figure 4*D*). Once the pool of reduced PC and cyt.*f* were exhausted, restricted PSII performance could have contributed to the lower *k*_end_ in anoxia.

It is of note that throughout the study, the partially complemented C1 line, which accumulates ∼75% of WT PGR5 levels under the control of its native promoter (19), resembled WT in oxic conditions whereas it tended to partially perform like *pgr5* in anoxia. With exception of P700 in anoxic conditions (Figure S4*B*), this was most apparent on the levels of *b*-hemes oxidation (*k*_*b*-ox_ in Figure 3*D*). For cyt.*f* reduction (*k*_*f*-red_ in Figure 3*C*) and electrogenicity (*k*_ini_ in Figure 4*C*), both rates were significant faster under anoxia as compared to *pgr5* but still slower than WT. To eliminate PGR5 titration effects in C1, we generated an independent PGR5-complemented line which, besides the expected P700 redox behaviour (Figure 5*A*), also produced WT-like cyt.*f* reduction rates after the pulse (*k*_*f*-red_ in Figure 5*B*), as well as *k*_*b*-ox_ (Figure 5*C*) and *k*_ini_ (Figure 5*D*). We noted that hetero-phototrophic cells varied slightly in their rates, compared to photoautotrophic cells (cf. Figures 1 and 2). As we will discuss later, the zeocin system showed that electrogenicity of the photosynthetic chain relied in part on the availability of the immediate P700 donor pool which was governed by Qi-site turnover in a PGR5-dependent manner. For feasible photoautotrophic culturing under low light, the C1 line with a native *PGR5* promotor is preferred over the zeocin-induced, complemented strain.

**Figure 5.**
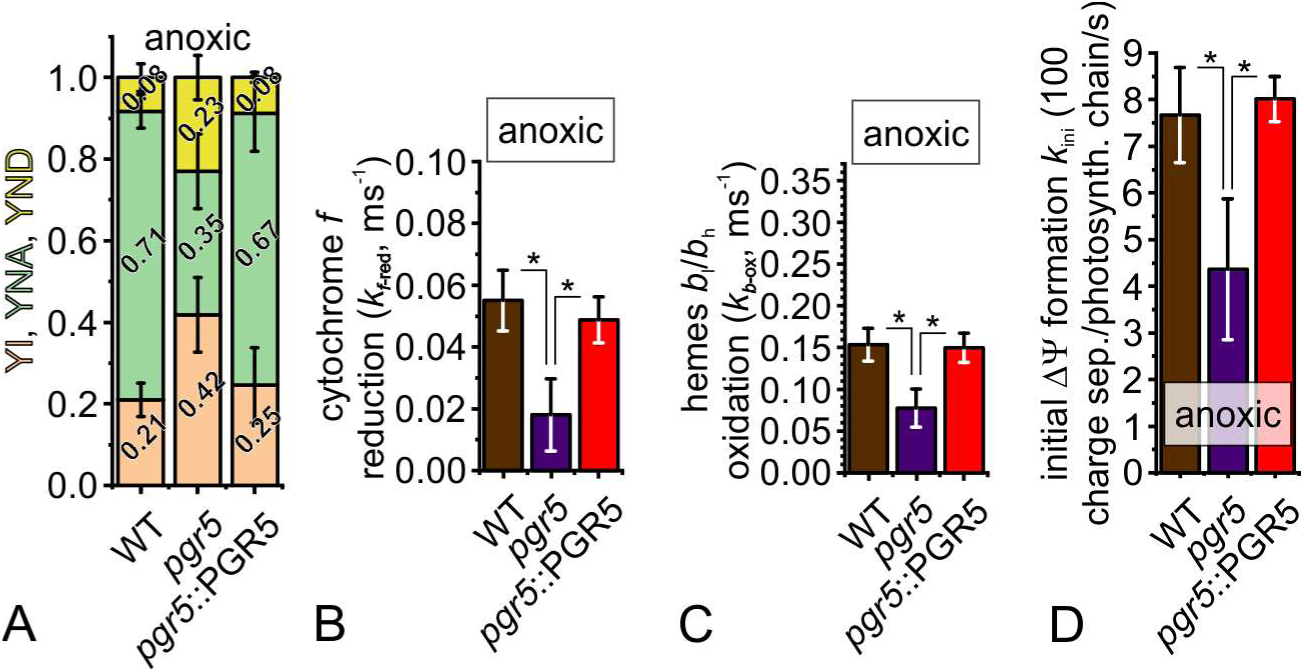
Assessments of steady state P700 redox state (A), cytochrome *b*_6_*f* dark redox relaxation (B, C), and electrogenic capacity of the photosynthetic chain (D) in anoxic WT (brown), *pgr5* (violet), and PGR5-complemented lines (red, *pgr5*::PGR5) show complementation of the *pgr5* phenotype in the rescued line. Cells were grown in Tris-acetate-phosphate medium and PGR5 expression was induced by 5 µg/mL zeocin 24 h before experiments. (A) The different P700 populations were deconvoluted as oxidisable fraction (YI, yield of PSI), non-oxidisable P700 owing to acceptor side limitation (YNA), and pre-oxidised fraction due to donor side limitation (YND). Like in Figure 2*B*, the anoxic *pgr5* maintained higher YI and YND, and failed to develop pronounced YNA like WT and *pgr5*::PGR5. (B) After the pulse, the reduction rates of cytochrome *f* (k_*f*-red_) were significantly slower in the mutant which recovered to WT-level in *pgr5*::PGR5. For kinetics refer to Figure S6*A*. (C) Simultaneously, the oxidation rates of *b*-hemes after the pulse (k_*b*-ox_) were significantly slowed down in the mutant only. Refer to Figure S6*B* for kinetics. (D) When steady state cells experienced the saturating pulse, the high initial electrogenic charge separation rates (*k*_ini_) in WT were PGR5-dependent (*N* = 3 ± SD; Welch’s t-test \**P* < 0.05).

### 3. PGR5-dependent redox finetuning of the *b*_6_*f* low-potential chain and implications on the Q cycle

In order to rule out possible dark redox equilibration artefacts (owing to different pre-oxidation levels in the steady state), this section introduces single turnover measurements. Here, the light-adapted cells have been assayed in absence of PSII photochemistry and upon a 30-s dark period. This setup ensures reduction of primary and secondary donors of PSI as well as *pmf* consumption, especially since the ΔpH governs photosynthetic control (14, reviewed in 15). The single *b*_6_*f* turnover passes the electron hole from oxidised *c*-heme in cyt.*f* to the Rieske ISP which, after swapping back the FeS domain closer to cytochrome *b*_6_ subunit, is reduced at the Qo-site in a bifurcated process that also reduces hemes *b*_l_/*b*_h_. When electrons are passed on heme-*b*_h_, a ΔΨ was generated which we monitored via ECS signals. Redox changes in the *b*_6_*f* (Figure 6*A*) and the corresponding ECS changes (Figure 6*B*) were assayed in oxic samples. The ECS kinetics in Figure 6 are relative and are composed of three phases (reviewed in 44, 46). First, the unresolved *a*-phase finished in less than 1 µs before the first measured signal and represented PSI photochemistry in this experiment. The *a*-phase upon the sub-saturating flash generated ∼0.4 – 0.5 charge separations per total PSI, serving for ECS kinetics normalisation. In the ∼10-ms range, the *b*-phase corresponded to the *b*_6_*f*-dependent ΔΨ generation and the *c*-phase was dominated by ATP synthesis activity which consumed the ΔΨ produced by the flash. In the ECS kinetics in Figure 6, the *b*-phase was deconvoluted by subtracting from the total ECS kinetics the *c*-phase, which instantaneously followed first-order exponential decay (47, 48). Regarding the decay rate of the *c*-phase, we observed no significant differences in the two strains (supplementary Figure S7). We also measured the redox kinetics and ECS in presence of MV (Figures 5*C* and 5*D*) as well as in anoxic conditions (Figures 5*E* and 5*F*). On a time scale after injecting an electron hole into the *b*_6_*f*, cyt.*f* reduction (*k*_*f*-red_ derived from single exponential fits in the *b*_6_*f* redox kinetics panels) preceded the electrogenic *b*-phase (*k*_ΔΨ_ derived from single exponential fits in the ECS kinetics panels) and the last phase was the relatively slow oxidation of hemes *b*_l_/*b*_h_ (*k*_*b*-ox_ derived from single exponential fits in the *b*_6_*f* redox kinetics panels). A statistical evaluation of *k*_*f*-red_ (Figure 6*G*), *k*_ΔΨ_ (Figure 6*H*), and *k*_*b*-ox_ (Figure 6*I*) is shown as well as the amplitude of the *b*-phase relative to one charge separations per PSI (Figure 6*J*).

**Figure 6.**
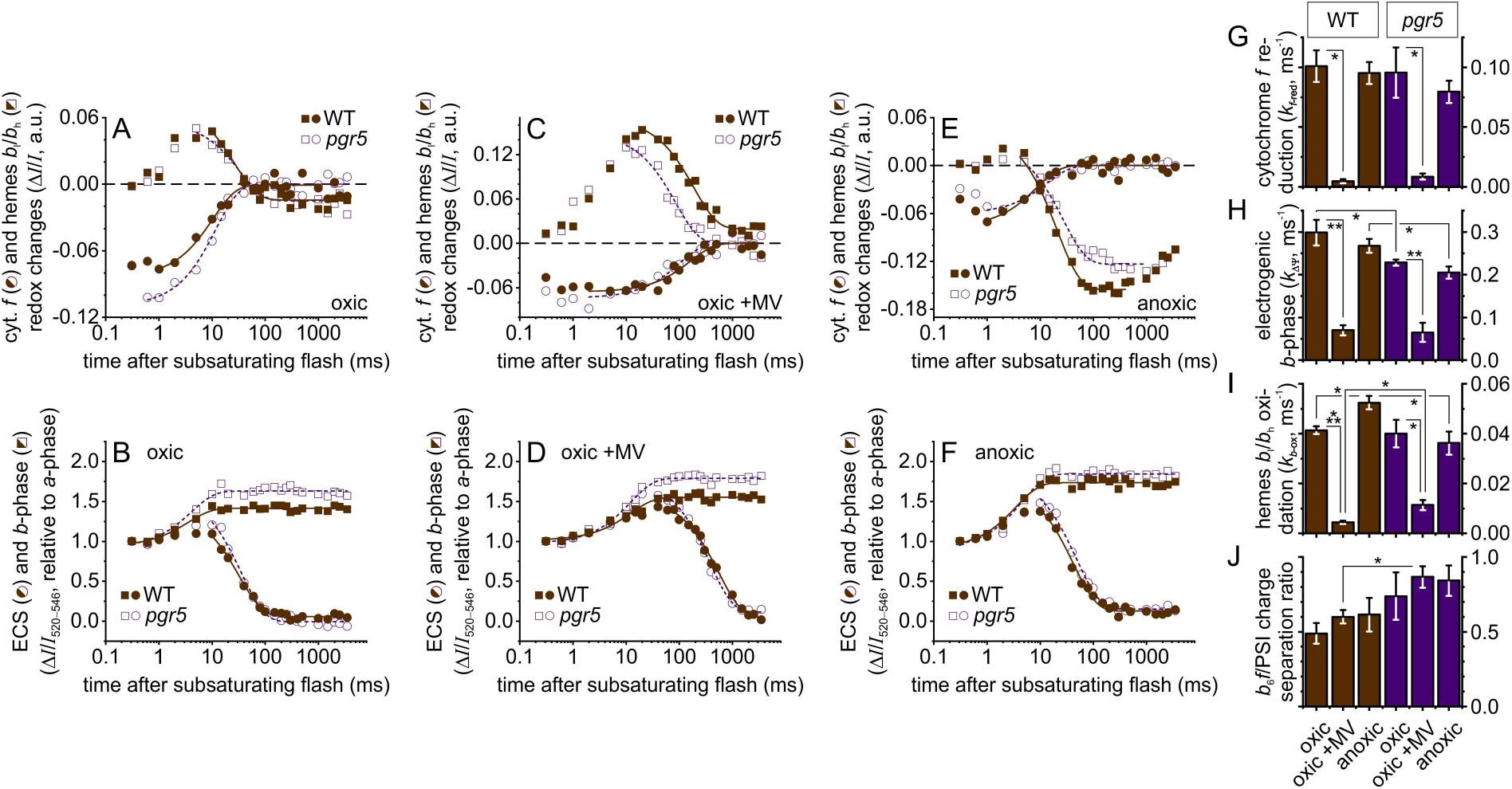
Redox kinetics and electrogenic signals reveal a PGR5-dependent low-potential chain tuning in anoxia as well as an inhibitory effect of MV on the single *b*_6_*f* turnover. (A) Cytochrome *f* and *b*-heme signals are shown for oxic WT and *pgr5*. Cytochrome *f* is rapidly (<300 µs) oxidised after the flash and re-reduced within 100-ms darkness (fitted curves). The *b*-heme net reduction lasted between ∼1-ms and 10-ms darkness, followed by a slower oxidation phase (fitted curves) for several tens of ms. (B) The corresponding oxic ECS kinetics were normalised to the signal produced by a flash hitting ∼40% of PSI centres (<300 µs, *a*-phase). The following *b*-phase (fitted curve), resulting from *b*_6_*f*-dependent charge separation activity in the ∼10-ms range, was deconvoluted from raw ECS kinetics by subtracting exponential decay phase produced by ATP synthase (*c*-phase, fitted curve, refer also to Figure S7). In this sample, the *b*-phase developed slower in *pgr5* and showed a slightly larger relative amplitude. (C) Addition of MV slowed down cytochrome *f* reduction and *b*-heme oxidation in both strains, and the mutant was slightly less affected. The inhibition of *b*-heme oxidation allowed larger reduction amplitudes compared to panel A. (D) Evolution of the *b*-phase was slowed down by MV but the amplitude was not altered. MV also slowed down the *c*-phase upon *γ*-disulfide promotion (72). (E) The *b*_6_*f* redox kinetics in anoxia showed slightly slower cytochrome *f* oxidation whereas reduction was like in oxic cells. Net reduction of *b*-hemes in the first 10-ms was of negligible amplitude and a large oxidation phase followed. (F) ECS and *b*-phase kinetics in anoxia resembled oxic panel B. (G) Cytochrome *f* reduction rates *k*_*f*-red_ were calculated from the fitted decay functions and showed significant slowdown in the presence of MV. (H) The electrogenic *b*-phase also evolved at slower rates (*k*_ΔΨ_) in MV samples. Compared to WT, the *pgr5* mutant showed slower *k*_ΔΨ_ in oxic and anoxic conditions. (I) After the apparent *b*-heme reduction phase ceased, the slow oxidation rates were expressed as *k*_*b*-ox_. MV slowed down *k*_*b*-ox_ and the WT showed faster *k*_*b*-ox_ in anoxia whereas the anoxic mutant *k*_*b*-ox_ remained unchanged. (J) The relative *b*-phase amplitudes, compared to the *a*-phase, were comparable and only larger in *pgr5* MV samples. The mutant had a tendency, though, to produce larger *b*-phase amplitudes. (*N* = 3 ± SD; Welch’s t-test \**P* < 0.05, \*\**P* < 0.005 and \*\*\**P* < 0.0005)

In oxic samples, independently of MV, cyt.*f* oxidation was finished before the first record at 300 µs after the flash and resulted in an amplitude of ∼-0.1 units compared to the reference signal before the flash (circle symbols in *b*_6_*f* redox kinetics panels). In anoxic samples, oxidation of cyt.*f* was slowed down slightly, finishing between ∼1-ms and ∼2-ms after the flash. With exception of MV samples, cyt.*f* reduction by the FeS domain was initiated at ∼1-ms, yielding similar *k*_*f*-red_ values in oxic and anoxic conditions. When MV was added, *k*_*f*-red_ was lowered significantly (5% and 9% residual rates of oxic WT and *pgr5*, respectively) and a delayed onset of reduction became apparent between ∼5-ms and ∼10-ms.

Before cyt.*f* was getting reduced in oxic and anoxic samples (during the first ms), net redox changes of hemes *b*_l_/*b*_h_ were very small (square symbols in *b*_6_*f* redox kinetics panels). Only after the onset of cyt.*f* reduction, a net reduction of hemes *b*_l_/*b*_h_ became apparent, which coincided with the *b*-phase (square symbols in ECS kinetics panels). The sequential reduction of cyt.*f* and hemes *b*_l_/*b*_h_ was expected and was also observed in MV samples but there was a significant slowdown of the low-potential chain turnover. The amplitude of hemes *b*_l_/*b*_h_ reduction in the presence of MV appeared larger since Qo-site turnover, with *k*_ΔΨ_ as proxy, was less slowed down than *k*_*b*-ox_ (23% and 26% residual *k*_ΔΨ_ of oxic WT and *pgr5* in Figure 6*H* vs. 10% and 28% residual *k*_*b*-ox_ in Figure 6*I*). The inhibitory MV effect on *k*_*b*-ox_ in *pgr5* was less efficient, which might account for smaller net reduction amplitude during the 10-ms phase (Figure 6*C*).

In both strains under oxic and anoxic conditions, the net reduction amplitudes in the hemes *b*_l_/*b*_h_ redox kinetics during the first 10-ms differed (Figure 6*A* and Figure 6*E*). During the initial phase in oxia and anoxia, the electron transfer rate *k*_ΔΨ_ between hemes *b*_l_ and *b*_h_ (which depends on *k*_*f*-red_ and the following electron injection into the low-potential chain upon Qo-turnover) was not changed in the respective strains. However, *k*_ΔΨ_ differed between WT and the less efficient *pgr5* (Figure 6*H*), although hemes *b*_l_/*b*_h_ net reduction amplitude was comparable (Figure 6*A* and Figure 6*E*). As discussed later, this suggests that oxidation of heme-*b*_h_ (yielding PQH_2_ eventually) was slower during the electrogenic 10-ms phase in *pgr5* and thereby allowed a similar net reduction amplitude compared to WT.

The relative amplitude of the *b*-phase was between 50-85% of the *a*-phase and tended to be slightly higher in *pgr5* (Figure 6*J*). Further experiments need to clarify whether less efficient PQH_2_ formation at the *pgr5* Qi-site, which would weaken stromal charge neutralisation compared to WT, was responsible for slightly higher *b*-phase amplitudes in the mutant. Nevertheless, the ΔΨ generated by one *b*_6_*f* turnover in our conditions was close to values in earlier reports which attributed similar fractions of one charge separation across the whole membrane when measuring electron transfer ‘within’ the membrane bilayer from hemes *b*_l_ to *b*_h_ in the *bc*_1_ complex (49, 50). After injection of Qo-site electrons into the low-potential chain, the slower hemes *b*_l_/*b*_h_ oxidation phase (*k*_*b*-ox_ in Figure 6*I*) produced faster rates in anoxic WT only. Whether the dysfunctional *k*_*b*-ox_ tuning was responsible for slower *k*_ΔΨ_ in *pgr5* needs to be examined.

## Discussion

PGR5 is an important regulator of photosynthetic electron transfer, however, its function has not been linked to the operation of the *b*_6_*f*. Our data indicate a dysfunctional *b*_6_*f* in the absence of PGR5. The *b*_6_*f* enigmatically receives a stromal feedback from PSI. It modifies the Q cycle in strongly reducing conditions or is inhibited when being disconnected from PSI signals in the presence of artificial electron acceptors. We provide evidence that PGR5 is functionally involved in a modified Q cycle which has access to stromal electrons and operates in WT but less efficiently in *pgr5*.

Please refer to supplementary Figure S8 in the supplementary discussion for a detailed interpretation of the results in moderately reduced stroma. Our *pgr5* findings under oxic conditions, arguing in support of previous reports (19, 34, 35), suggest unregulated coordination of electrons between PSI and its sinks that produced higher PSI-borne LEF rates and resulted in a more oxidised electron carrier pool downstream of PSII.

We did also observe a more oxidised electron carrier pool in anoxic *pgr5* – but under these reducing steady state conditions we located it downstream of the *b*_6_*f*. The undersupply of electron donors rendered PSI more oxidisable by the pulse and thus diminished acceptor side limitation in *pgr5* (cf. *A* and *B* panels of Figures 1, 2, and Figure 5*A*). Accordingly, the high-potential chain was even more oxidised in the steady state (cf. cyt.*f* oxidisability in *C* panel inserts of Figures 1, 2, and Figure S6*A*) and redox relaxation in the mutant *b*_6_*f* was slowed down in the dark (Figures 3*C* and 3*D*, Figures 5*B* and 5*C*). The PGR5-dependent bottleneck at *b*_6_*f* exhausted the immediate PSI donor pool and thus lowered the initial ΔΨ formation rate significantly when rapidly transitioning anoxic mutant cells to strong light (*k*_ini_ in Figures 4*C* and 5*D*). Notably in the single turnover measurements (Figure 6), which were carried out in absence of PSII activity when ΔpH was collapsed and photosynthetic control was cancelled, the *pgr5* mutant was generally less efficient to separate charges across the membrane via the *b*_6_*f* (*k*_ΔΨ_ in Figure 6*H*) and failed to accelerate the electron discharge from the low-potential chain in anoxia (*k*_*b*-ox_ in Figure 6*I*). This argues that the absence of PGR5 has a direct impact on *b*_6_*f* function.

The observed steady state *pgr5* phenotype in anoxic conditions can also be linked to the *b*_6_*f* function when the WT showed enhanced Qi-site activity upon *k*_*b*-ox_ redox tuning (Figure 7*A*). This tuning feature prevents deleterious electron back-up in the low-potential chain, which we will explore further below, and thus promotes photosynthetic (self-)control via the *b*_6_*f* in high light conditions. It is of note that light acclimation in the cuvette (i.e., steady state actinic light) did not increase photosynthetic control since we did not observe increased YND in the WT (Figure 1*B*). However, *k1*_*P*-red_ in anoxic cells was slowed down after the saturating pulse (Figure 3*B*), suggesting enhanced photosynthetic control compared to oxic conditions. Importantly, YND (established before the pulse) and *k1*_*P*-red_ (measured after saturating pulse) were probed at two different *pmf* levels. A closer look at the electric *pmf* component after the pulse shows that the ΔΨ was consumed back to the steady state level in about 10-ms to 20-ms darkness (cf. insert of Figure 4*A*). Therefore, it could be that photosynthetic control was established in the course of the pulse and contributed to *k1*_*P*-red_.

**Figure 7.**
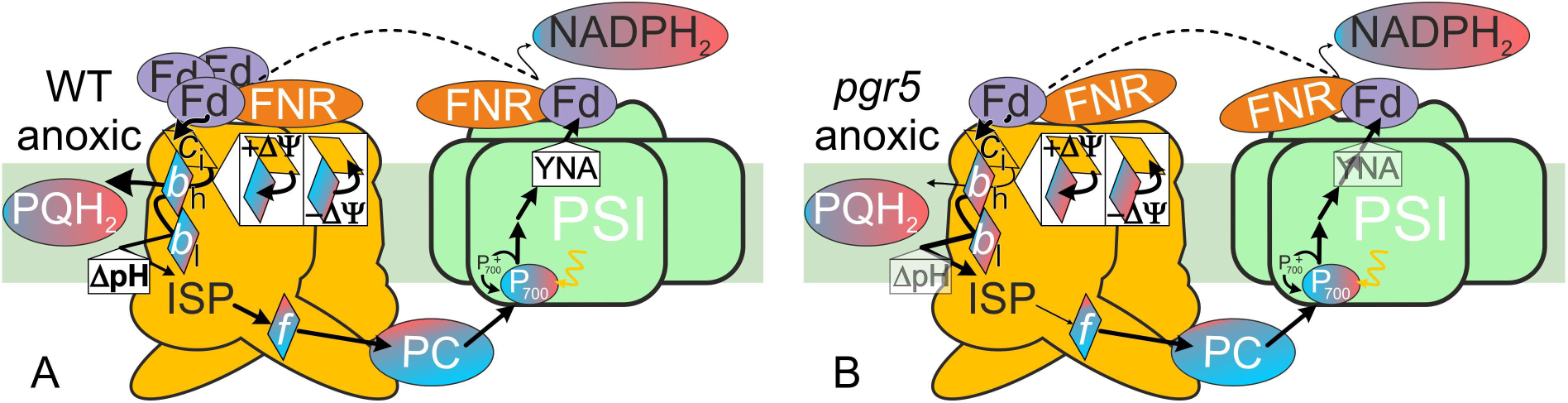
Model summarizing the multiple-turnover measurements under anoxic conditions. Except for PC and NADPH_2_, the redox levels were measured (blue and red stand for oxidized and reduced, respectively). PSI acceptor side limitation (YNA) and photosynthetic control (ΔpH) were established between weakly (transparent) and strongly contributing levels (bold). Modified forward reaction efficiencies are highlighted (Qo: PQH_2_ oxidation, Rieske ISP/cyt.*f* interaction; Qi: PQ reduction, *b*_h_/*c*_i_ electron sharing, Fd-dependent *c*_i_ reduction; NADPH_2_ formation). Refer to Figures 8 and S1 for details on *b*_h_/*c*_i_ interaction and stromal electron utilization. (A) When anoxic WT *b*_6_*f* operates in the Fd-assisted Q cycle mode, *b*_6_*f* electrogenicity is maintained and ΔpH protects PSI. (B) Inefficient utilisation of excess stromal electrons at the Qi-site stalls the *pgr5 b*_6_*f* in anoxic conditions, thus weakening YNA. The effect could be linked to aggravated, incomplete FNR tethering (20) and a disturbed low-potential chain oxidation influences ISP/cyt.*f* interaction (12). Refer to Figure S8*C* for a similar scenario in the presence of MV.

In anoxic conditions, the *pgr5* mutant failed to discharge low-potential chain electrons (for explanation, see scheme in Figure 7*B*). Manifested in single turnover experiments, the *k*_*b*-ox_ redox tuning defect eventually impaired the steady state turnover. The inhibited steady state *b*_6_*f* in anoxic *pgr5* would produce less YNA due to increased YND, as observed. The low-potential chain electron back-up in the steady state might slowdown the *b*-heme re-reduction phase in anoxic *pgr5* (*k*_*b*-red_ in Figure 3*E*) once these electrons were discharged upon ΔΨ consumption in the dark. Given the significant *b*-heme oxidation (*D* panels of Figures 1, 2, and S4) and the ΔΨ-dominated *pmf* when probing the ECS decay amplitude in the dark (Figures 4*A*, 4*B*, and S5), electron back-up in the low-potential chain was probably also contributing to the poor steady state *b*_6_*f* turnover in MV samples (supplementary Figure S8*C*). Interpretation of the steady state samples can be inferred from the single turnover experiments in the presence of MV where, in both cases, *k*_*b*-ox_ was significantly slower (Figure 3*D*, and Figure 6*I*). The inhibition of *k*_*b*-ox_ by MV suggests that *b*_6_*f* function relies on stromal redox signals associated with PSI turnover.

Figure 7 summarises the multiple-turnover measurements under anoxic conditions in a scheme and proposes a safety mechanism of the *b*_6_*f* turnover (see respective arrow heads in Figure 7), which couples *b*-heme oxidation to cyt.*f* reduction via the Rieske ISP (reviewed in 41, 51-53). In the following, we will draw the attention to the *b*_6_*f* turnover and focus on the low-potential chain which, as introduced, underlies a canonical Q cycle. Thus, the *b*-heme kinetics under oxic conditions in Figure 6*A* followed the transient net reduction/oxidation and, compared to the pre-flash reference signal, the slightly more oxidised *b*-hemes after ∼100 ms resulted from a small pre-reduced heme-*b*_h_ population (Figure S1). Despite the drawback that both *b*-hemes were spectroscopically monitored as one population at room temperature (42) and we could not determine the heme-*c*_i_ redox state, our observed changes in the low-potential chain oxidation behaviour under anoxic conditions (Figure 6*E*) support a modified Q cycle with access to stromal electrons (see below).

However, the initial fast oxidation phase (≤ ∼10-ms after the flash, see two left panels in Figures 7*A* and 7*B*) can be explained on grounds of a canonical Q cycle since the samples were strongly pre-reduced. The 30-s darkness before the flash were sufficient to pre-reduce heme-*b*_h_ but not heme-*b*_l_ (42, 43), and to reduce heme-*c*_i_ in absence of ΔΨ (2). The heme-*b*_h_ dark reduction rates in anoxic samples (42, 43) indicate that *b*_h_^red^/*c*_i_^red^ was very likely to occur, yielding the pre-reduced Qi-site. Yet, pre-reduction was not sufficient to drive PQH_2_ formation in the dark since the necessary ΔΨ prerequisite was not met (3) and *b*_h_^red^ oxidation was only achieved with the flash. Regarding the probed *b*-hemes, the anoxic samples contained *b*_l_^ox^/*b*_h_^red^ before the flash, and *b*_l_^ox^/*b*_h_^ox^ at the end of the measurement (thus the negative *b*-hemes signals in Figure 6*E*). The relatively flat *b*-heme redox signals during 10-ms were a result of *b*_l_^red^/*b*_h_^red^ disqualification (9, 10, 11 and references therein) so that the fast *b*_h_^red^ oxidation (at a rate ≤ *k*ΔΨ) was driven by reduction of *b*_l_^ox^ upon Qo-site turnover. To explain the faster oxidation phase in WT vs. *pgr5* after the electrogenic 10-ms phase (*k*_*b*-ox_ in Figure 6*I*), a modification of the canonical Q cycle is required. With ongoing consumption of ΔΨ, *b*_h_^red^/*c*_i_^ox^ would transiently convert to *b*_h_^ox^/*c*_i_^red^ (2), which eventually makes the Qi-site ‘PQ-accessible’ for the next turnover upon weakening the ligation of *c*_i_^red^ with F40 in subunit-IV (7). We propose that this canonical scenario produced the corresponding *k*_*b*-ox_ in anoxic *pgr5* (see two right panels in Figure 8*A*). In anoxic WT, after ∼10-ms darkness, *b*_h_^red^/*c*_i_^ox^ has access to stromal electrons so that *b*_h_^red^/*c*_i_^red^ is formed and a second PQH_2_ is produced at the Qi-site (see two right panels in Figure 8*B*). The accessible substrate in the WT Qi-site allows faster *b*_h_^red^ oxidation and is not governed by the ΔΨ-dependent transition to *b*_h_^ox^/*c*_i_^red^, unlike in the mutant (cf. similar ECS decay in Figures 5*F*, and S7). The second oxidation of *b*_h_^red^ (yielding *k*_*b*-ox_ beyond ∼10-ms) is slower than the one during the first Qi-site turnover due to absence of the reducing pressure on the *b*_6_*f* (Qo-site electrons entering the low-potential chain). A facilitated generation of *c*_i_^red^ from stromal electrons could increase the chance for the, compared to oxic conditions, less abundant PQ to enter the Qi-site binding pocket (7). By enhancing low-potential chain oxidation, this WT *b*_6_*f* feature is an optimisation to highly reducing conditions, as evidenced by our steady state observations where the anoxic *pgr5* displayed an arrested *b*_6_*f*. In reducing conditions, the mutant *b*_6_*f* underlies the intrinsic, short-circuit-preventing process that influences cyt.*f* reduction rate by governing the interaction between the Rieske ISP FeS domain and cytochrome *b*_6_/cyt.*f* (12). Unless *b*-hemes are not oxidised in steady state *pgr5*, the reduced flexible FeS domain will not swap closer to cyt.*f* to release the “trapped” high-potential chain electron (reviewed in 41).

**Figure 8.**
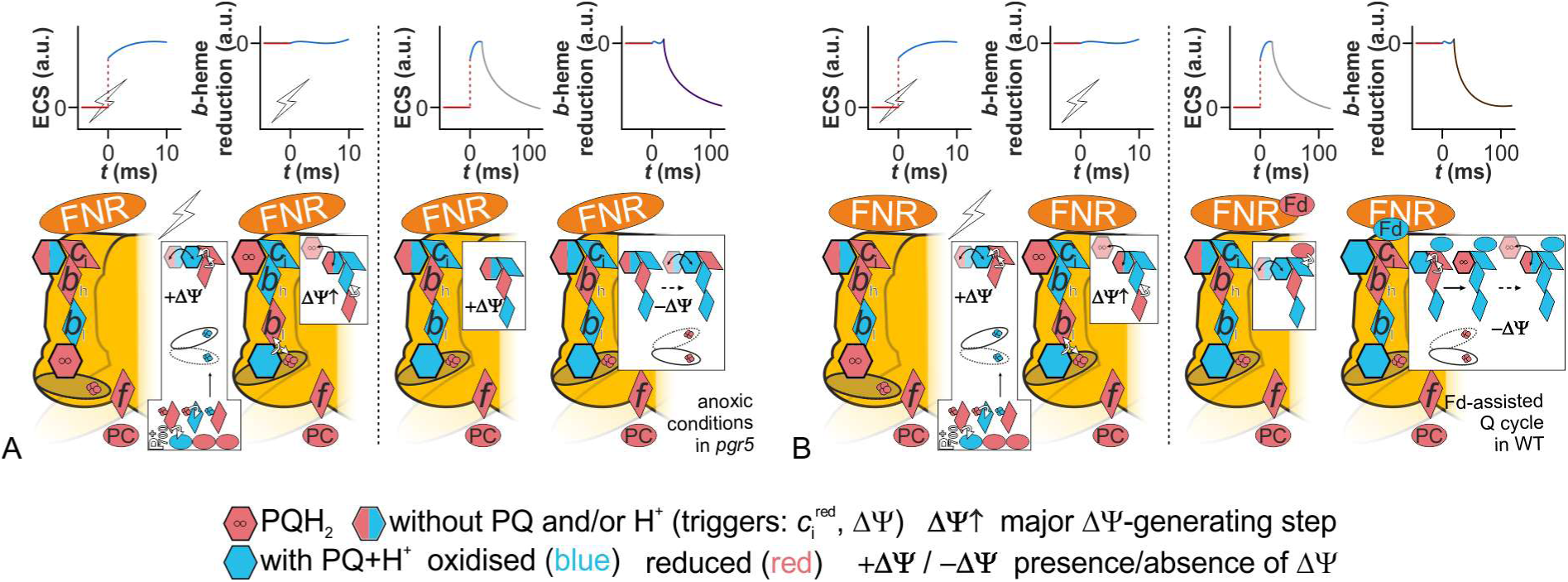
A proposed sequence of *b*_6_*f* redox reactions in anoxic algae highlights the differences in the *pgr5* mutant (A) compared to WT (B). Schematics of the electrochromic shift (ECS) and the *b*-heme net reduction are shown for the corresponding time windows. The initial reaction steps were very similar in both strains (left of the dashed line in panels A and B). The dark-equilibrated pre-flash *b*_l_^ox^/*b*_h_^red^ re-equilibrated again to *b*_l_^ox^/*b*_h_^red^ after the Qo-site turnover and electrogenic charge transfer (unaltered *b*-heme net reduction). During this initial phase, one PQH_2_ is formed at the Qi-site by making use of the pre-reduced *b*_h_^red^/*c*_i_^red^ redox couple. Although not indicated here, this reaction is slightly slower in *pgr5* when taking slower *b*_6_*f* electrogenicity as a proxy (cf. Figure 6*H*). (A) The following *b*-heme oxidation phase is slower in the mutant and may be attributed to poor Qi-site substrate availability which is promoted upon *c*_i_^red^ formation (7). In presence of a membrane potential (ΔΨ), the *b*_h_^red^/*c*_i_^ox^ redox couple equilibrates (2) and a relatively slow transition to *b*_h_^ox^/*c*_i_^red^ follows the ΔΨ decay in *pgr5* (grey and violet curve). (B) Unlike the mutant, WT retains sufficiently more FNR to the thylakoid membrane (20) which might be an allosteric modulator by interacting with the *b*_6_*f* (23-25). Thus, Fd bound to FNR might drive the transient generation of *c*_i_^red^ in the presence of ΔΨ, which then creates the ‘PQ-accessible’ Qi-site specifically in WT and, due to the *b*_h_^red^/*c*_i_^red^ redox couple, produces faster *b*-heme oxidation. Compared to *pgr5*, the faster *b*-heme oxidation also accelerates the Rieske FeS domain to swap closer to cytochrome *f* (reviewed in 41), which could be important during multiple turnover operation as indicated in Figure 7.

In conclusion, PGR5 is involved in the Q cycle operation under anoxic conditions which attributes Fd-PQ-reductase activity to the *b*_6_*f*, as originally suggested by Arnon (54). A stromal redox poise activates this specific *b*_6_*f* function, which increases the H^+^/e ratio of 2 that is obtained in LEF-favouring conditions during a canonical Q cycle. Under CEF-promoting conditions, the role of PGR5 is to ensure reduction of the heme-*c*_i_ in the *b*_6_*f*, likely via Fd. The *b*_6_*f* low-potential chain activity in *pgr5* is slightly lower during single turnovers and steady state activity in CEF-favouring conditions requires the PGR5 polypeptide. Thus, PGR5 is required for efficient Q cycle in the *b*_6_*f*, which is in turn crucial for lumen acidification to trigger the onset of photosynthetic control and qE. Whether PGR5 action during the redox poise is direct or indirect needs to be further investigated. In yeast-based protein-protein interaction studies, candidates for PGR5 interaction were cytochrome *b*_6_ and Fd (34). The PGR5-dependent CEF effector proteins PETO and ANR1, which interact with the algal *b*_6_*f* (55-58), are part of the Fd interactome as well (59, 60) and could have Fd shuttling functions. The yeast two-hybrid study also showed that PGRL1 interacts with PGR5 and FNR (34), and BiFC studies indicated protein-protein interaction between PGRL1 and ANR1 (57). Notably, FNR has been shown to co-purify in *b*_6_*f* preparations (23-25) and the algal *pgr5* retains less FNR on the thylakoid membrane (20). Interestingly, MV-treated thylakoid membranes do not retain FNR either (61). Thus, FNR membrane recruitment itself might (in)directly rely on reduced Fd since it is stimulated in anoxic conditions and requires functional *b*_6_*f* and PSI (62). By disturbing the electron flow downstream of PSI, probably on the level of Fd (34, 35), the FNR-anchoring trigger might be weakened in *pgr5*. This may lower functional Fd availability to reduce the heme-*c*_i_, whereas other sinks like hydrogenase might successfully compete for Fd in anoxic *pgr5* (63, 64). One could imagine that bound FNR could have allosteric effects on Qi-site turnover since the *b*_6_*f* redox kinetics in the presence of MV, in which FNR is expected to be soluble (61), showed attenuated resemblance of *b*_6_*f* samples treated with the Qi-site inhibitors MOA-stilbene (10) and NQNO (65). Further experiments are required to provide mechanistic insights into the PGR5-dependent electron transfer, permitting reduction of the *b*_6_*f* Qi-site during alternative Q cycle.

## Materials and methods

### Strains and cell cultures

As described previously (19), *Chlamydomonas reinhardtii* WT strain t222+, *pgr5* and a complemented line, termed C1, were used. The complemented C1 strain accumulated ∼75% of WT PGR5 levels (19). Cells were cultivated on agar-supplemented plates in Tris-acetate-phosphate medium (TAP, 66) at 20 µmol photons m^-2^ s^-1^. When growing cells for experiments, liquid Tris-phosphate medium was devoid of acetate (TP). Stirred cultures were grown at 10 µmol photons m^-2^ s^-1^ (16 h light/8 h dark) and were bubbled with sterile air at 25 °C. Grown cultures were diluted ∼6-fold at least once after inoculation and grown to a density of ∼2 × 10^5^ cells mL^-1^ before harvesting (5000 rpm, 5 min, 25°C). For experiments with PGR5-complemented lines that express zeocin resistance, WT, *pgr5* and *pgr5*::PGR5 were grown in TAP in same conditions as for TP cells, except without air bubbling. One day before the experiments, cells were diluted in fresh TAP and 5 µg/mL zeocin was added to *pgr5*::PGR5 cultures. Cells were resuspended at 20 µg chlorophyll mL^-1^ in TP supplemented with 20% (*w*/*v*) Ficoll and shaken vigorously in dim light. Optionally for oxygen-deprived conditions in the dark, cells were supplemented with 50 mM glucose, 10 U glucose oxidase and 30 U catalase in a cuvette, and then overlaid with mineral oil for at least 30 min before measurements. In presence and absence of PSII photochemistry, these illuminated cells will be referred to as anoxic.

### Generation of PGR5 complemented lines using dicistronic system

The *PGR5* gene (Cre05.g242400) was amplified from genomic DNA extracts using forward (5’-GCCCC<u>GAATTC</u>ATGCTGGCCTCCAAGCCCGTTGTTG) and reverse oligos (5’-CTAG<u>TCTAGA</u>TTAAGCCAGGAAGCCAAG) that harboured the underlined *Eco*RI/*Xba*I restriction sites. The digested fragment was introduced into a dicistronic expression vector (67, 68) for PGR5 under the control of the *PSAD* promoter. The construct conferred zeocin resistance as well, since *PGR5* expression was linked to *ble* via a skipping peptide FMDV2A, and DNA was introduced to the *pgr5* nuclear genome by electroporation (25 µF, 1kV). Transformants were pre-selected on TAP agar plates supplemented with 10 µg/mL zeocin.

### Chlorophyll fluorescence analysis

The LED-based spectrophotometer (JTS-10, BioLogic, France) was equipped with a Fluo_59 accessory in fluorescence mode. Probing was performed with 520-nm LED measuring pulses (350 µmol photons m^-2^ s^-1^) and light-detecting photodiodes were protected from scattered actinic light by using appropriate 3-mm thick filters (reference diode: BG39; measuring diode: LPF650+RG665, Schott, Mainz, Germany). After 30-min dark-adaptation, F_0_ was detected by probing in the dark and F_v_/F_m_ ((F_m_−F_0_)/F_m_) was obtained by a 250-ms saturating pulse (520-nm LED, 5000 µmol photons m^-2^ s^-1^). Adaptation to 630-nm actinic light (LEDs emitting 4700 µmol photons m^-2^ s^-1^ which yielded ∼150 µmol photons m^-2^ s^-1^ at the measuring cuvette) was carried out by regularly resuspending the sample for at least 30-min in an open cuvette. Oxygen-deprived cells were not mixed. The photochemical quantum yield of PSII, Φ_PSII_, was calculated from (F_m_’−F_s_)/F_m_’ by probing maximal (F_m_’) and steady state (F_s_) fluorescence in the light. The PSII efficiency factor, *qP*, was derived from (Φ_PSII_×F_m_)/F_v_ (69). The Q_A_^−^ re-oxidation in the dark was calculated as described elsewhere by using the Stern-Volmer relationship (32). When assuming a lake model for light harvesting, the redox state of the PQ pool in the light, 1−*qL*, was estimated from (F_m_’−F_s_)/(F_m_’−F_0_’)×F_0_’/F_s_ (40) with F_0_’ being derived from F_0_/(F_v_/F_m_+F_0_/F_m_’) (70).

Picked colonies, putatively carrying the dicistronic *PGR5* construct, were grown at 10 µmol photons m^-2^ s^-1^ and submitted to spot tests performed on TP agar plates (supplementary Figure S9). Selection was based on WT-like chlorophyll fluorescence after 24-h exposure to 200 µmol photons m^-2^ s^-1^, using a Maxi-Imaging PAM chlorophyll fluorometer (Walz, Germany).

### Time-resolved absorption spectroscopy

Time-resolved measurements are expressed as ΔI/I and were carried out with the JTS-10. P700 redox changes were measured with combined detection LEDs at 705 and 740 nm and associated interference filters at 705 and 740 nm (FWHM: 6 nm and 10 nm respectively). The light-detecting diodes were protected from scattered actinic light by a RG695 filter (Schott, Mainz, Germany). For reference purpose, PSII inhibitors hydroxylamine (HA at 1 mM final, from 1M aqueous stock) and 3-(3,4-dichlorophenyl)-1,1-dimethylurea (DCMU at 10 µM final, from 10 mM ethanolic stock) were added to obtain maximal P700 oxidation during the saturating pulse. However, the kinetics shown in the paper contained light-adapted cells (at least 30-min with 630-nm LEDs emitting 4700 µmol photons m^-2^ s^-1^) which had a functional PSII. The continuous actinic red light was hatched by short dark intervals (250 µs) during which 10-µs detecting pulses were placed after 200 µs. The P700 redox kinetics was recorded during saturating pulse (12-ms of 630-nm LED, 3000 µmol photons m^-2^ s^-1^) and followed for several seconds darkness. The P700 kinetics in the dark were corrected for a linear drift that had developed in light-adapted cells especially in anoxic cells (supplementary Figure S10). ECS signals were measured as the difference of the absorbance changes at 520 and 546 nm (respective interference filters FWHM: 10 nm) using white pulsed LED probing light. The light-detecting diodes were protected from scattered actinic light by 3-mm BG39 filters (Schott, Mainz, Germany). A saturating pulse (22-ms of 630-nm LED, 3000 µmol photons m^-2^ s^-1^) was used for kinetics of ECS and the cytochrome *b*_*6*_*f* redox reactions. Using appropriate interference filters (FWHM: 10 nm), the latter were monitored on the level of cytochrome *f* (554 nm) and cytochrome *b* (563 nm) with a baseline drawn between 546 and 573 nm (71). At variance (71), 554-nm signals were corrected with 0.23 × (563 nm – 546 nm) to subtract the contribution of cytochrome *b* to cytochrome *f* kinetics (4), especially in reducing conditions where this correction yielded pre-flash cytochrome *f* signals after several tens of ms (supplementary Figure S11). In single-turnover kinetics, the *b*-phase (electrogenic *b*_*6*_*f* contribution to the ECS signal) was deconvoluted by subtracting the *c*-phase (ATP synthase activity that results in ECS decay) which followed first-order exponential decay (47, 48). All ΔI/I signals in this study were calibrated to ECS changes produced by one charge separation upon a saturating laser flash in the presence of PSII inhibitors HA and DCMU. The PSI:PSII ratios were obtained by referring to uninhibited flash ECS amplitudes which were produced upon two charge separations by both photosystems, as reviewed recently (44). The overall membrane potential formation in saturating light was determined via a Dark Interval Relaxation Kinetics-based protocol (45), using ECS signal slopes at the end of the 22-ms light pulse and in subsequent darkness. Dark signals were disregarded during the first 2-ms, since they might be prone electrogenic PSI charge recombinations (38), and slopes up to 16-ms were calculated with the linear fitting function in OriginPro software (OriginLab). The same fitting was used for ECS slopes during the first 2-ms upon changing the light intensity. To normalise to one charge separation, the ECS slope values in uninhibited samples were divided by the flash-induced ECS ΔI/I in presence of HA/DCMU (charge separations/photosynthetic chain/s). Unless otherwise stated for calculated decay rates, kinetics were fit with mono-exponential decay function “ExpDec1” in OriginPro software and the fast P700 component, coined *k1*_*P*-red_ in the text, was obtained via two-exponential fitting “ExpDec2” (dark kinetics from 4-ms to 2550-ms). For *k*_*f*-red_ and *k*_*b*-ox_ the 600-ms phase was used for ExpDec1 fitting. However, a shorter time window for anoxic *k*_*b*-ox_ calculation was necessary for a seamless transition to the ExpDec1 fit of *k*_*b*-red_ up to 9-s dark.

## Acknowledgments

M.H. acknowledges support from Deutsche Forschungsgemeinschaft (DFG) Grant HI 739/13-1 and Grant HI 739/13-2

