## Supplemental Information for "PGR5 is required for efficient Q cycle in the cytochrome *b*_6_*f* complex during cyclic electron flow"

Table of contents


- Supplementary Table S1
- Supplementary Figures S1 – S11
- Supplementary Discussion
- Supplementary References

Supplementary Table S1: Maximal PSII quantum yield, Fv/Fm, of dark-adapted cells in different cellular redox states, as well as the PQ pool redox status, 1-*qL*, upon various light treatments.

| **Sample (*N*=3 ± SD)** | **Fv/Fm (D)** | **light treatment** | **1-*qL*** |
|---|---|---|---|
| wt oxic | 0.69 (±0.01) | D +10-s AL | 0.69 (±0.04) |
| | | AL 30 min | 0.65 (±0.06) |
| | | AL + MV | 0.72 (±0.01) |
| wt anoxic | 0.57 (±0.02) | D +10-s AL | 0.92 (±0.03) |
| | | AL 30 min | 0.83 (±0.00) |
| *pgr5* oxic | 0.70 (±0.03) | D +10-s AL | 0.67 (±0.02) |
| | | AL 30 min | 0.57 (±0.03) |
| | | AL + MV | 0.70 (±0.05) |
| *pgr5* anoxic | 0.58 (±0.02) | D +10-s AL | 0.91 (±0.04) |
| | | AL 30 min | 0.91 (±0.02) |
| C1 oxic | 0.69 (±0.01) | D +10-s AL | 0.66 (±0.01) |
| | | AL 30 min | 0.66 (±0.00) |
| | | AL + MV | 0.71 (±0.02) |
| C1 anoxic | 0.60 (±0.03) | D +10-s AL | 0.94 (±0.01) |
| | | AL 30 min | 0.89 (±0.02) |

D… dark-adapted; AL… actinic light with LEDs emitting 4700 µmol photons $m^{-2}$ $s^{-1}$, corresponding to ~150 µmol photons $m^{-2}$ $s^{-1}$ at the measuring cuvette; MV… 10 mM methyl viologen

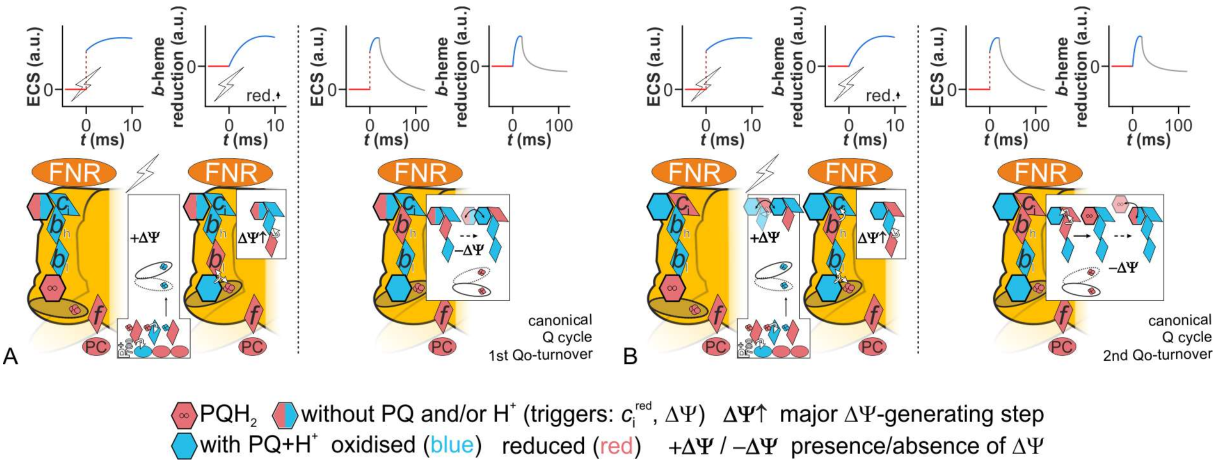


Supplementary Figure S1. Operation of the $b_6f$ via a canonical Q cycle is shown, including schematics of the developing electrochromic shift (ECS) signals and *b*-heme redox changes. Two consecutive Qo-site turnovers are shown, each induced by a laser flash. (A) The ellipse symbolizes the mobile FeS domain of the Rieske ISP, which swaps to a position close to the cytochrome $b_6$ subunit after reducing cytochrome *f*. After reduction of the FeS domain by $PQH_2$, heme-$b_l$ reduces heme-$b_h$ which generates membrane potential ($\Delta\Psi$). With decay of the $\Delta\Psi$, the low-potential chain redox carrier heme-$c_i$ gets reduced by heme-$b_h$ (1). Reduction of heme-$c_i$ opens the Qi-site for the substrate quinol (2), which ligates with heme-$c_i$. Swapping of the FeS domain closer to the high-potential chain cytochrome has been linked to *b*-heme oxidation (reviewed in 3). (B) In the presence of $\Delta\Psi$ upon the flash, heme-$c_i$ reduces heme-$b_h$ (1). The oxidised heme-$c_i$ ligates more tightly with substrate quinol which then requires more reducing pressure on the low-potential chain to inject the first electron into the ligated ensemble (2). This energy is provided by the oxidant-induced reduction of heme-$b_l$ which drives, simultaneously, oxidation of heme-$b_h$ since a fully reduced *b*-heme population is has not been observed during single turnover measurements (4, 5, 6 and references therein). Presence of a $\Delta\Psi$ is important for oxidation of the low-potential chain (7) and the (semi-)PQ in the Qi-site receives the electrons from $c_i^{red}$ and/or $b_h^{red}$ probably in a concerted and closely spaced process.

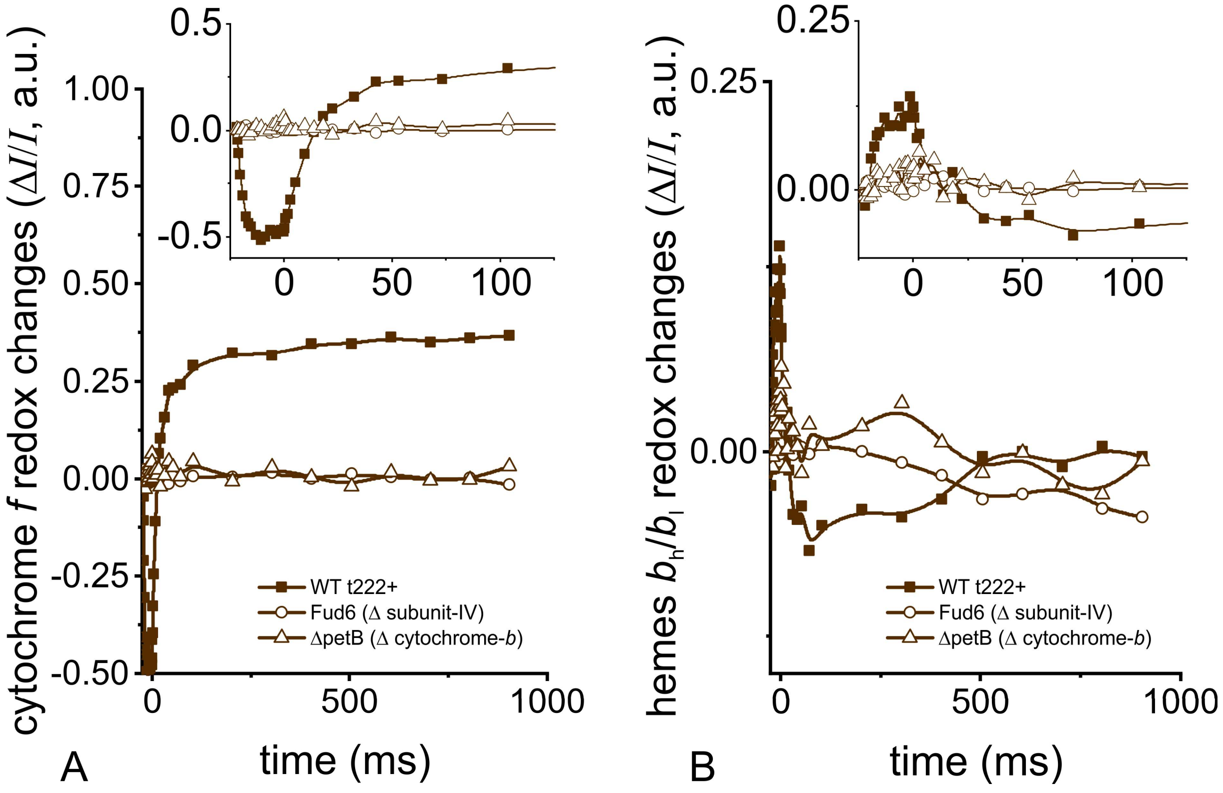


Supplementary Figure S2. The measured redox signals of cytochrome *f* (A) and *b*-hemes (B) were assayed in $b_6f$-lacking strains Fud6 (8, 9) and ΔpetB (10). During the pulse (insets) the respective signal amplitudes were small in the mutants. Cells were grown in Tris-acetate-phosphate medium in dim light.

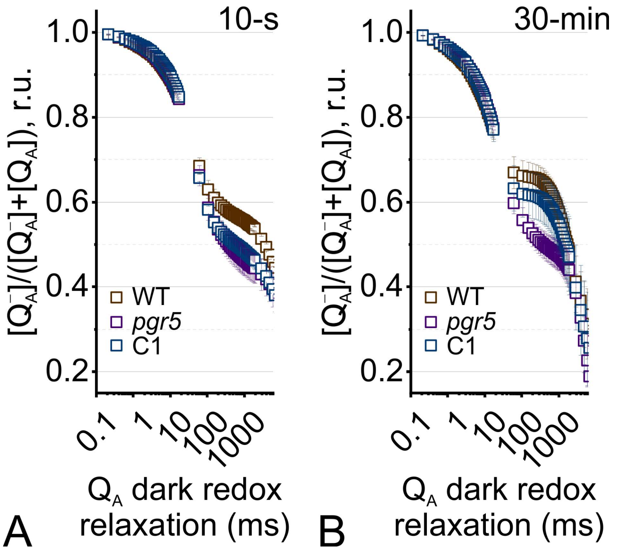


Supplementary Figure S3. Following a saturating-pulse in oxic cells, redox relaxation of $Q_A^-$ in the dark are shown (*N* = 3 ± SD). (A) Kinetics after short illumination of dark-adapted cells for 10-s and (B) kinetics after light adaptation for 30-min are shown. The fully reduced $Q_A^-$ pool was re-oxidised in samples by ~30% in the first ~50-ms. Only in panel B the retardation phase up to ~500-ms was present, except for *pgr5*.

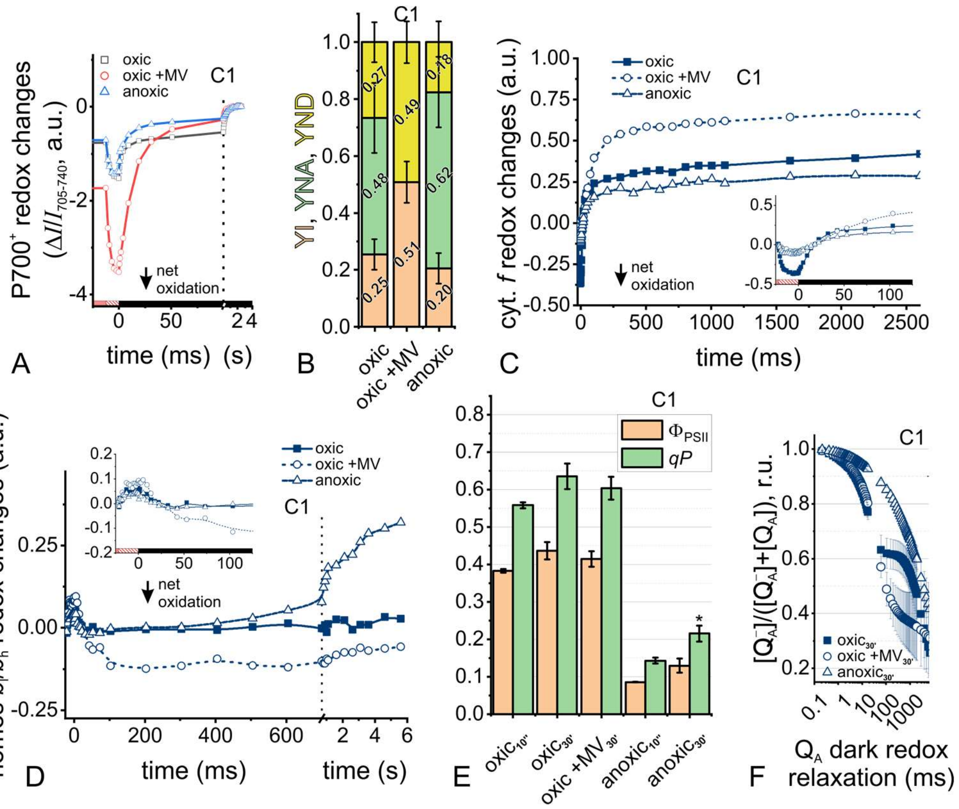


Supplementary Figure S4. Total electron transfer from PSII to PSI is changed in the C1 line under different cellular redox states. (A) Saturating pulse-induced P700 redox changes in absence and presence of 10 mM methyl viologen (MV) are shown, as well as in anoxic cells. The 12-ms pulse (hatched red box) was applied on light-adapted cells in the steady state (red box), followed by several seconds dark measurements (black box). (B) The different P700 populations were deconvoluted as oxidisable fraction (YI, yield of PSI), non-oxidisable P700 owing to acceptor side limitation (YNA), and pre-oxidised fraction due to donor side limitation (YND). The electron acceptor MV abolished YNA and increased YND despite PSII activity. Anoxic conditions lowered YI and increased YNA. (C) Cytochrome *f* redox changes in a similar light/pulse/dark regime show different pulse-induced oxidation magnitudes (inset) with oxic > anoxic ~ oxic+MV amplitudes, compared to the steady state reference. MV addition caused no significant cytochrome *f* net oxidation during the pulse. Most of the dark relaxation finished in ~50-ms, except for slow re-reduction in MV samples. (D) Redox changes of *b*-hemes were comparable during the pulse (inset) and most of the oxidation was finished in ~50-ms, ~300-ms, ~50-ms in oxic, oxic +MV and anoxic samples, respectively. The latter samples showed a slow re-reduction phase with an onset at less than 300-ms dark. (E) Chlorophyll fluorescence-derived quantum yield of PSII ($\Phi_{PSII}$) and PSII efficiency factor (*qP*) are shown ($N = 3 \pm$ SD). Anoxia lowered *qP* and $\Phi_{PSII}$ compared to oxic conditions, and *qP* was slightly higher than in anoxic$_{30'}$ WT (Welch's t-test $P < 0.05$, cf. Figure 1 of the main text). (F) Following a saturating-pulse in light-adapted cells, redox relaxation of $Q_A^-$ in the dark are shown ($N = 3 \pm$ SD). The fully reduced $Q_A^-$ pool was re-oxidised in oxic samples by ~30% in the first ~50-ms and a following ~500-ms retardation phase was missing upon MV addition. $Q_A^-$ re-oxidation was slowed down in anoxic cells.

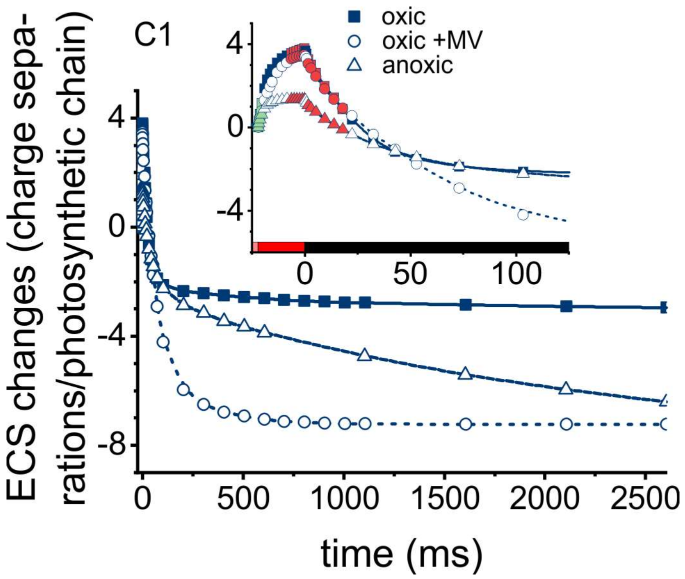


Supplementary Figure S5. The ability of the C1 line to generate and dissipate an electric field is shown by measuring the electrochromic shift, ECS. Signals of samples from Figure S4 were recorded in steady state light, during a saturating pulse and in darkness. The kinetics indicate that the 22-ms pulse led to equilibration of a new membrane potential ΔΨ. The efficiency to generate a higher ΔΨ level at the onset of the pulse is shown by initial rates in green (inset), yielding $k_{\text{ini}}$ from the linear slope during the first 2-ms of the pulse. The ΔΨ generation capacity at the end of the pulse ($k_{\text{end}}$) was corrected with the ΔΨ-consuming activity by ATP synthase. Values are shown in panels *C* and *D* of Figure 4 in the main text.

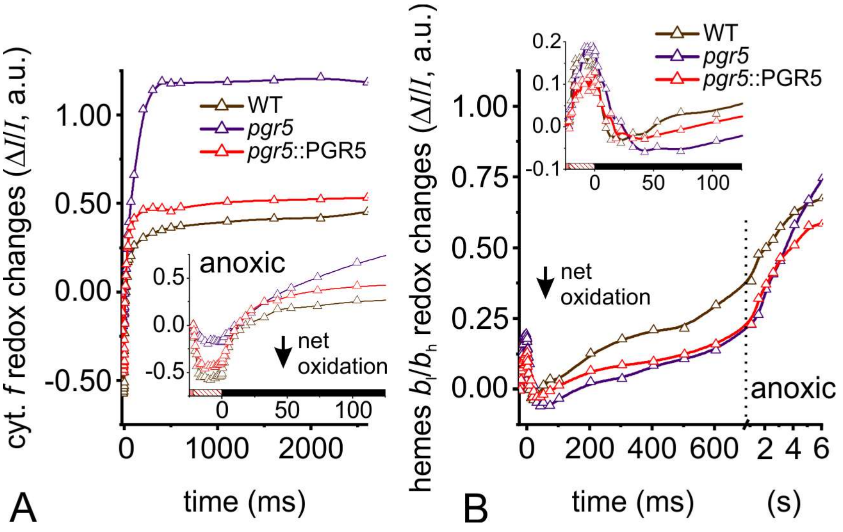


Supplementary Figure S6. The measured redox signals of cytochrome *f* (A) and *b*-hemes (B) were assayed in anoxic WT (brown), *pgr5* (violet) and a complemented *pgr5* line harbouring a complementation construct (red). The latter was a dicistronic vector that conferred selection for zeocin resistance and put *PGR5* expression under the control of a *PSAD* promoter (11, 12). Cells were grown in Tris-acetate-phosphate medium in dim light. (A) Cytochrome *f* redox changes in a light/pulse/dark regime show different pulse-induced oxidation magnitudes (inset) with WT ~ *pgr5*::PGR5 > *pgr5*, compared to the steady state reference. Refer to Figure 5*B* in the main text for calculated cytochrome *f* reduction rates after the pulse. (B) Redox changes of *b*-hemes were comparable during the pulse (inset) and most of the oxidation was finished in ~25-ms, ~50-ms, ~25-ms in WT, *pgr5* and *pgr5*::PGR5, respectively. Refer to Figure 5*C* in the main text for calculated *b*-hemes oxidation rates after the pulse.

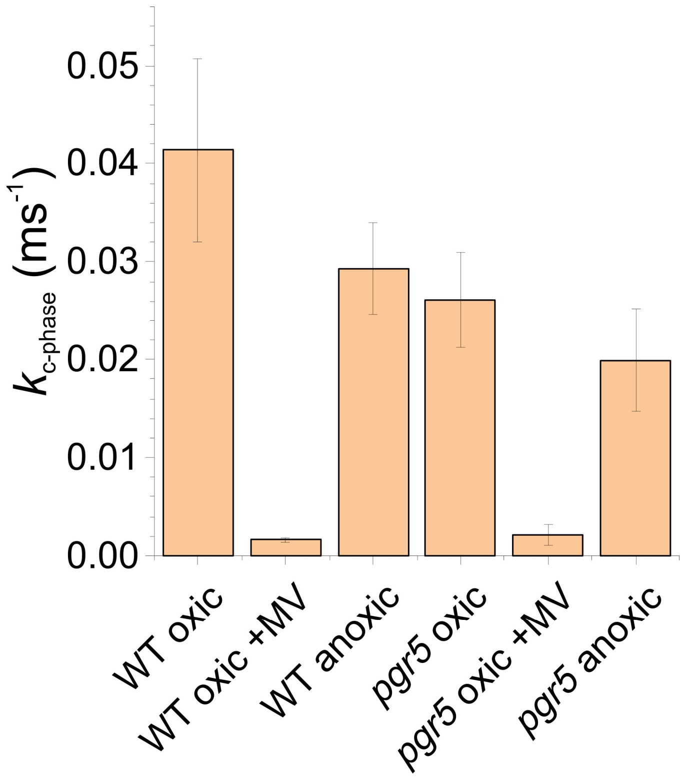


Supplementary Figure S7. The exponential ECS decay rates of the *c*-phase, $k_{\text{c-phase}}$, was produced by ATP synthase activity. Please do refer to ECS panels in Figure 6 of the main text. In the respective conditions, WT and *pgr5* did not produce significantly different $k_{\text{c-phase}}$ ($N = 3 \pm$ SD).

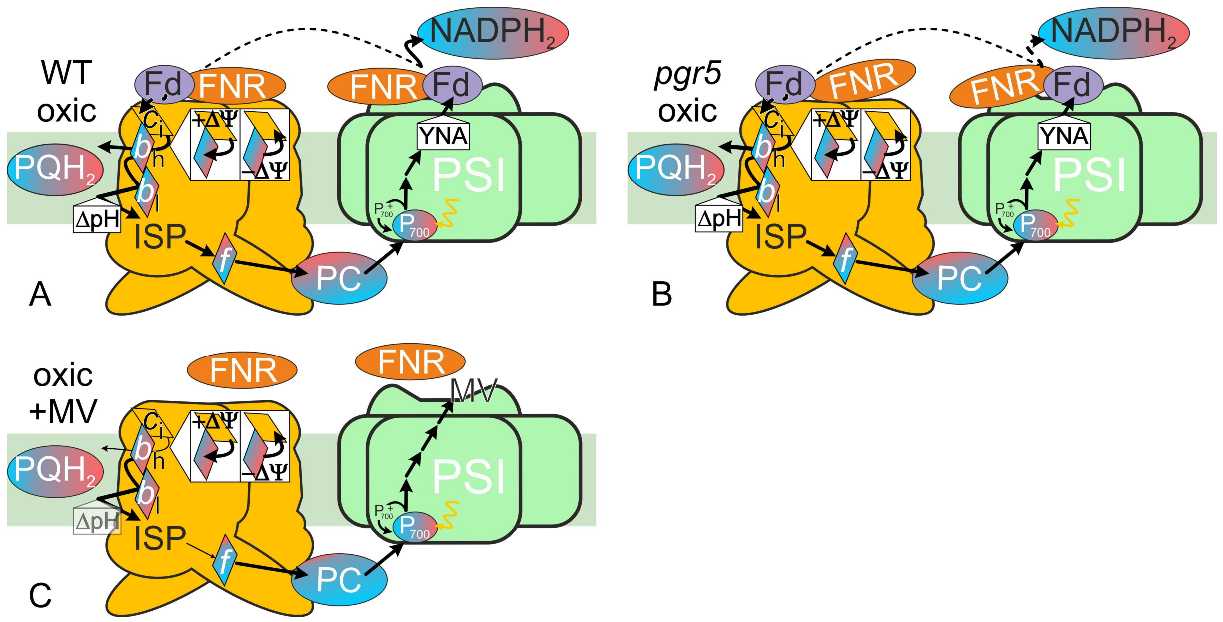


Supplementary Figure S8. Model summarizing the multiple-turnover measurements under oxic conditions. Except for PC and $NADPH_2$, the redox levels were measured (blue and red stand for oxidized and reduced, respectively). PSI acceptor side limitation (YNA) and photosynthetic control ($\Delta pH$) were established, optionally absent or at weakly contributing levels (transparent). Modified forward reaction efficiencies are highlighted (Qo: $PQH_2$ oxidation, Rieske ISP/cyt.*f* interaction; Qi: PQ reduction, $b_h/c_i$ electron sharing, Fd-dependent $c_i$ reduction; $NADPH_2$ formation). Refer to Figure S1 for details on $b_h/c_i$ interaction. Unlike oxic WT (A), *pgr5* (B) shows unregulated electron utilisation downstream of PSI (dashed arrow). Since Fd pool was not reduced in the experiments it could sustain faster electron flow via PSI without producing YNA. Still, the PQ pool and cyt.*f* were slightly more oxidised in the mutant. FNR membrane recruitment is impaired in the mutant (13). (C) Qi-site inhibition by methyl viologen (MV) disturbs low-potential chain oxidation and, thus (14), ISP/cyt.*f* interaction. MV likely interferes with the redox signal that mediates FNR tethering to the membrane (15, 16).

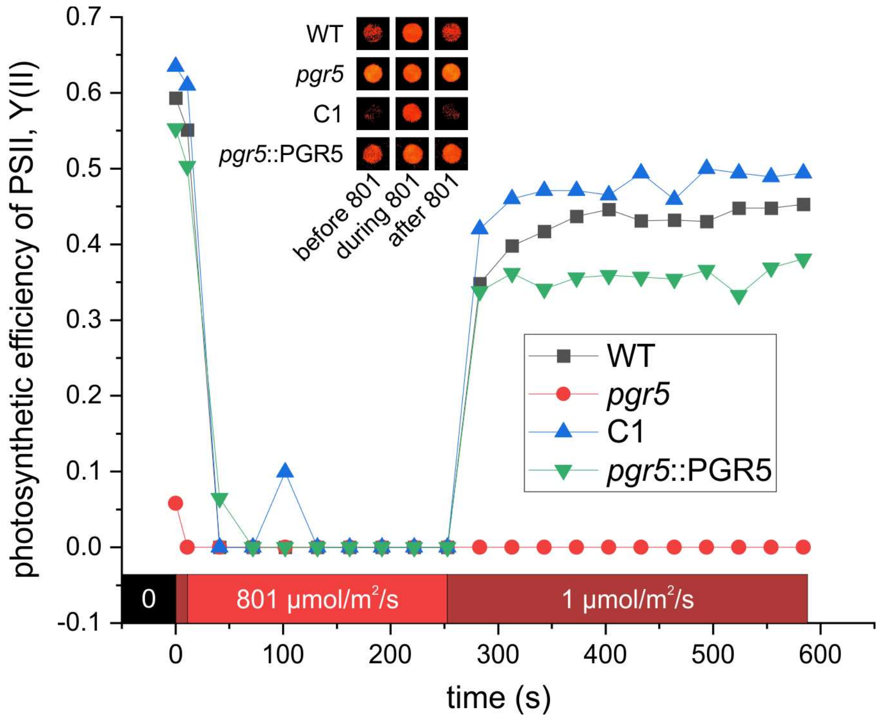


Supplementary Figure S9. Spot test of a PGR5-complemented line (*pgr5*::PGR5; grown on plates in the presence of 10 µg/mL zeocin) facilitated transformant selection based on chlorophyll fluorescence. Measurements were carried out with cells after 24-h exposure to 200 µmol photons/m$^2$/s, using a Maxi-Imaging PAM chlorophyll fluorometer (Walz, Germany). WT, *pgr5*, and C1 were grown in absence of zeocin and only *pgr5* showed low Y(II). The inset shows exemplary chlorophyll fluorescence records of the spots before, during and after strong actinic light, corresponding to 801 µmol photons/m$^2$/s.

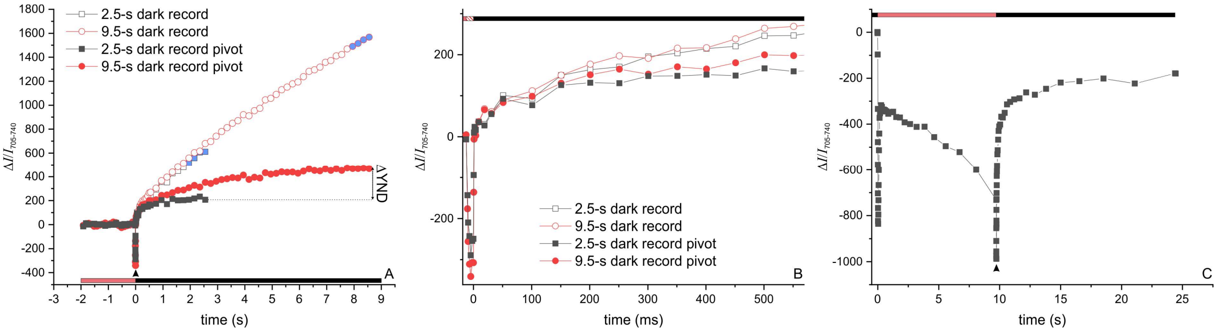


Supplementary Figure S10. Light-adapted anoxic cells displayed a linear drift after illumination. (A) ΔI/I 705-740 kinetics of light-adapted cells are shown and P700 net oxidation resulted in negative signals. Kinetics were recorded in the light (red bar) and, after a saturating pulse (arrowhead), for different durations in the dark (black bar). Open symbols are kinetics with the drift in the dark and blue symbols were the linear regression correction for the partial kinetics after the pulse. The corrected traces are shown as filled symbols. Depending on the dark interval, the corrected kinetics differed in their fractional donor side limitation (ΔYND) which translated in altered YNA as well. The photo-oxidisable fraction was not affected since the deconvolution of YI was made with data recorded in the light, and thus not subjected to correction. (B) Close-up of the fast P700 re-reduction phase. The drift correction had marginal consequences on the apparent fast (≤ 100-ms) P700 re-reduction rate. (C) Illumination of anoxic cells for 10-s did not produce a dark drift and majority of P700 re-reduction was finished in the 2.5-s time window, which we used to drift-correct the kinetics in the main text.

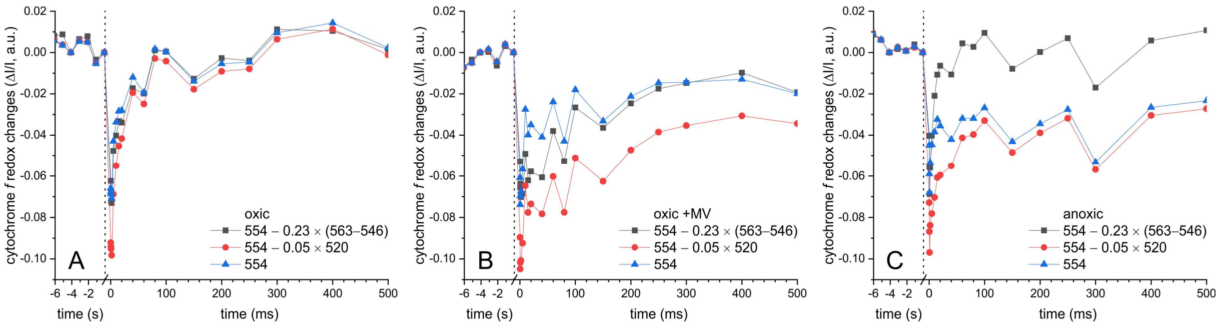


Supplementary Figure S11. Deconvolution of the cytochrome *f* signals required a correction for partial *b*-heme contributions. Oxic cells in the absence (A) and presence of MV (B) are shown, as well as anoxic samples (C). The cytochrome *f* redox signals were monitored at 554 nm with a baseline drawn between 546 and 573 nm (17). These curves are shown as blue triangles. Previous reports (17) subtracted 5% of the 520-nm signal, using the same baseline, to correct for ECS contributions (red circles). We noticed especially in anoxic samples that pre-flash levels ($x \leq 0$) were neither obtained from 554 nm signal deconvolution nor in the ECS-corrected kinetics. Instead, by estimating extinction coefficients cytochrome *b* at 554 nm (18), the cytochrome *f* signals at 554 nm were corrected by subtracting 0.23 × (563 nm – 546 nm). Thereby, the contribution of cytochrome *b* to cytochrome *f* kinetics was cancelled and pre-flash signals were obtained (black squares). This deconvolution was used in the main text. The ΔI/I signal in this figure was normalised to the ECS (520 nm – 546 nm) *a*-phase amplitude of active PSI.

Supplementary Discussion

Here we wish to focus on oxic conditions (moderately reducing stroma) which had, compared to anoxic samples, a different impact on the malfunctioning photosynthetic apparatus that contributed to the *pgr5* phenotype. When the PSI acceptor pool is oxidised, the *pgr5* results argue for higher LEF rates since we observed a more oxidised electron carrier pool downstream of PSII (higher *qP* and $\Phi_{PSII}$ in steady state *pgr5*, see in main text *E* panels of Figures 1, 2, and Figure S4), thus confirming earlier findings (19). These observations might be linked to a respiratory overflow valve in the algal *pgr5* mutant that resulted from a malfunctioning regulation of stromal redox homeostasis (19), and/or it might depend on flavodiiron proteins which are efficient electron sinks by reducing oxygen under these conditions in Chlamydomonas (20). Thus, respiration and/or flavidiiron proteins might maintain WT-like YI in oxic *pgr5*, grown and analysed in the permissive light conditions. On the contrary, Arabidopsis *pgr5* shows well-documented PSI overreduction (21). This phenotype can be alleviated in the *pgr5* background upon expression of flavodiiron proteins (22), which are absent in angiosperms. Higher *pgr5* LEF rates might be PSI-borne since faster P700 oxidation rates (main text Figure 3*A*) could produce lower cyt.*f* oxidisability by a strong pulse (cf. negative signal amplitude in main text *C* panel inserts of Figures 1, 2, and Figure S4), after equilibrating with the oxidized PC pool in steady state *pgr5*. Accordingly, the slightly lower electrogenic competence in oxic *pgr5* ($k_{ini}$ in main text Figure 4*C*) could be linked to electron exhaustion in the PC pool. This would prevent PSI from efficiently separating multiple charges during the turnovers in the 2-ms range. When measuring the transiently slowed down $Q_A^-$ re-oxidation, *pgr5* failed to establish this feature which was exclusive to light-adapted WT and C1 cells (Figure S3). The $Q_A^-$ re-oxidation kinetics up to ~500-ms darkness can be interpreted as a signature of the $b_6f$-mediated equilibration of PQ and PC pools (23), shuttling more electrons to the latter in oxic *pgr5* after the onset of darkness. Notably, the $Q_A^-$ re-oxidation signature was light adaptation-dependent and probably involves PGR5-dependent accumulation of stromal electrons downstream of PSI. Unlike in the WT (Figure S8*A*), the electron flow in *pgr5* was unregulated downstream of PSI when the Fd pool was not reduced yet (Figure S8*B*). In our growth and experimental conditions, PSI photochemistry was not inhibited like in high-light cultures (19), nor was the PSI:PSII ratio affected (13). Therefore, YNA was not increased in the mutant (Figure S8*B*), which one would expect at a lower PSI:$b_6f$ ratio where more electrons are funnelled through the remaining PSI centres (19). The $b_6f$ underlies an intrinsic, short-circuit-preventing process that influences cyt.*f* reduction rate by governing the interaction between the Rieske ISP FeS domain and cytochrome $b_6$/cyt.*f* (14). Unless *b*-hemes are not oxidised, the reduced flexible FeS domain will not swap closer to cyt.*f* to release the "trapped" high-potential chain electron (reviewed in 3). The electrogenic $b_6f$ defect in mutant single turnover measurements ($k_{\Delta\Psi}$) could result from electron back-up due to lower Qi-site activity. According to the above-mentioned coupling of $PQH_2$ formation at Qi to cyt.*f* reduction, a slowdown of high-potential chain electron supply would be expected in the steady state. However, the $b_6f$ in oxic *pgr5* could re-reduce P700 like WT and dark-relaxed internally in a comparable manner on the levels of cyt.*f* and hemes $b_l$/$b_h$. This suggests suppression of the $k_{\Delta\Psi}$ effect when $\Delta\Psi$ was present. Unblocking a Qi-site defect under continuous illumination has been observed before in an algal $b_6f$ point mutant (2). It remains enigmatic by which mechanism the $b_6f$ "senses" the $\Delta\Psi$, which is crucial for hemes $b_l$/$b_h$ oxidation (7).
